# Engineering functional human gastrointestinal organoid tissues using the three primary germ layers separately derived from pluripotent stem cells

**DOI:** 10.1101/2021.07.15.452523

**Authors:** Alexandra K. Eicher, Daniel O. Kechele, Nambirajan Sundaram, H. Matthew Berns, Holly M. Poling, Lauren E. Haines, J. Guillermo Sanchez, Keishi Kishimoto, Mansa Krishnamurthy, Lu Han, Aaron M. Zorn, Michael A. Helmrath, James M. Wells

**Affiliations:** College of Medicine, University of Cincinnati, Cincinnati, OH, 45267, USA; Center for Stem Cell and Organoid Medicine (CuSTOM), Cincinnati Children’s Hospital Medical Center (CCHMC), Cincinnati, OH, 45229, USA; Divisions of Developmental Biology, Cincinnati Children’s Hospital Medical Center (CCHMC), Cincinnati, OH, 45229, USA; Divisions of Pediatric General and Thoracic Surgery, Cincinnati Children’s Hospital Medical Center (CCHMC), Cincinnati, OH, 45229, USA; Endocrinology, Cincinnati Children’s Hospital Medical Center (CCHMC), Cincinnati, OH, 45229, USA; CuSTOM-RIKEN BDR Collaborative Laboratory, CCHMC, Cincinnati, OH, 45229, USA; Laboratory for Lung Development, RIKEN Center for Biosystems Dynamics Research (BDR), Kobe, 650-0047, Japan

**Keywords:** tissue engineering, gastric, enteric nervous system, mesenchyme, human pluripotent stem cells, Brunner’s glands

## Abstract

The development of human organoid model systems has provided new avenues for patient-specific clinical care and disease modeling. However, all organoid systems are missing important cell types that, in the embryo, get incorporated into organ tissues during development. Based on the concept of how embryonic organs are assembled, we developed an organoid assembly approach starting with cells from the three primary germ layers; enteric neuroglial, mesenchymal, and epithelial precursors, all separately derived from human pluripotent stem cells. From these we generated human gastric tissue containing differentiated glands, surrounded by layers of smooth muscle containing functional enteric neurons that controlled contractions of the engineered tissue. We used this highly tractable system to identify essential roles for the enteric nervous system in the growth and regional identity of the gastric epithelium and mesenchyme and for glandular morphogenesis of the antral stomach. This approach of starting with separately-derived germ layer components was applied to building more complex fundic and esophageal tissue, suggesting this as a new paradigm for tissue engineering.

## INTRODUCTION

All organs of the gastrointestinal (GI) tract are assembled from cells derived from the three primary germ layers during embryonic development. These diverse cell types are required for the proper execution of the GI tract’s complex functions. For example, key functions of the stomach to chemically and mechanically breakdown orally ingested nutrients depend on a complex interaction of the epithelium to produce acid and proteases, the smooth muscle to contract and relax, and the enteric nerves to provide input and coordinate both of these processes (Eicher, Berns and Wells, 2018). These three main components of the stomach develop separately from the three primary germ layers with the endoderm forming the epithelial lining, the mesoderm contributing to the stromal cells and smooth muscle layers, and the ectoderm giving rise to the enteric nervous system (ENS), yet come together to form a complete and complex layered structure that is functional by the time of birth. Then, each germ layer plays essential roles in postnatal function. The gastric ENS, as an intrinsic postganglionic network of excitatory and inhibitory neurons, along with the vagus nerve, coordinates the epithelial release of acid and proteases (Furness *et al*., 2020; Zhao *et al*., 2008; Norlen *et al*., 2005; Rydning *et al*., 2002) and the relaxation of smooth muscle needed for gastric emptying (Sung *et al*., 2018; Beckett, Sanders and Ward, 2017; Shaylor *et al*., 2016; Li *et al*., 2011).

During organ development, the ENS, mesenchyme, and epithelium communicate with each other in a temporally dynamic manner to regulate regional identity, morphogenesis, and differentiation of progenitors into specific cell types (Le Guen *et al*., 2015). For example, neural progenitors of the ENS, enteric neural crest cells (ENCCs), actively migrate to the foregut tube in response to signals from the surrounding mesenchyme at the same time as this mesenchyme is differentiating into multiple, organized layers of smooth muscle. Work with chick embryos have identified specific signaling pathways that mediate reciprocal signaling between germ layers. One common reciprocal signaling module involves sonic hedgehog (Shh), which is secreted by the epithelium of several developing organ and regulates expression of bone morphogenetic protein (BMP) in adjacent mesenchyme. BMP activation then mediates secondary responses such as patterning the developing gut tube mesenchyme (Roberts *et al*., 1995; Roberts *et al*., 1998; Faure *et al*., 2002; De Santa Barbara *et al*., 2005) and inducing epithelial cell fates like the pyloric sphincter at the junctions of developing organs (Smith and Tabin, 1999; Smith *et al*., 2000; Moniot *et al*., 2004; Theodosiou and Tabin, 2005). Epithelial Shh is also known to indirectly regulate ENCC proliferation and differentiation through manipulation of key extracellular matrix proteins within the gut mesenchyme (Nagy *et al*., 2016). Additional studies in chick embryos have shown that ENCCs are required for and regulate the growth, pattering, and maturation of developing stomach mesenchyme (Faure *et al*., 2015). Finally, recent work using both chick and mouse embryos have also shown that both epithelial Shh-induced BMP signaling and ENCC-produced BMP antagonists work in spatiotemporal concert to regulate the radial position of the gut’s smooth muscle layers (Huycke *et al*., 2019).

Congenital disorders arising from improper ENS and smooth muscle development include neuropathies that can impact the function of the proximal GI tract (Westfal and Goldstein, 2017). This results in dysregulation of motility and gastroesophageal reflux disease, collectively described as abnormal gastric emptying. A much more common gastric dysfunction that develops postnatally is gastroparesis. This involves an inability of the pyloric sphincter to relax in coordination with gastric contraction, preventing gastric contents from exiting the stomach which causes bloating and nausea. While the cause of this disorder is unknown, recent work using mouse embryos suggest it may be the result of hypoganglionosis or inflammatory degradation of intrinsic nNOS-expressing inhibitory neurons (Baker *et al*., 2020; Westfal and Goldstein, 2017). Surprisingly little is known about gastric ENS development in any model system, and development of a functional human gastric model system could accelerate not only the study of environmental and genetic factors impacting gastric motility, but also the discoveries of new therapies to improve gastric function.

While animal models have been invaluable to study stomach development and disease (de Santa Barbara, van den Brink and Roberts, 2002), there are major structural and functional differences in this organ between different species. For example, rodents have a forestomach that does not exist in humans. The avian stomach contains a proventriculus, which is a proximal glandular compartment, somewhat paralleling the human fundus, and a gizzard, which is a more distal grinding compartment that is vastly different than the human antrum (Kim and Shivdasani, 2016). There are also developmental differences at the molecular level; Hedgehog signaling appears to play opposite roles in the development of GI smooth muscle between chick and mouse embryos (Huycke *et al*., 2019). Each of the existing model systems have unique experimental strengths and weaknesses to study how the stomach forms from the three separate germ layers. Chick embryos are easy to manipulate *in vitro*, but are not a good genetic model. In contrast, mice are a strong genetic model, but germ layer specific studies *in vivo* are technically demanding or impossible.

We reasoned that by recapitulating organ assembly from the three germ layers *in vitro* we could both construct a more complex organoid and study human organ development in a way not previously possible. In this study, we incorporated human pluripotent stem cell (hPSC)-derived splanchnic mesenchyme and ENCCs into developing human antral and fundic gastric organoids (hAGO and hFGO, respectively) to recapitulate normal gastric development. The resulting gastric organoids were composed of epithelial glands surrounded by multiple layers of functionally innervated smooth muscle. The technology was readily transferrable to other types of organoids and was used to engineer esophageal organoids containing all three germ layers. The tractability of this approach allowed us to study germ layer communication during stomach development. We found that human ENCCs promote gastric mesenchymal development and glandular morphogenesis and that the presence of adequate amount of gastric mesenchyme is essential for maintaining gastric regional identity.

## RESULTS

### Deriving mesenchyme from hPSCs and incorporation into gastric organoids

One of the first and most critical steps in GI development is the assembly of epithelium and mesenchyme into a primitive gut tube. Establishing this basic epithelial-mesenchymal structure is essential for all subsequent stages of GI development. While PSC-derived human gastric organoids have a full complement of epithelial cell types (McCracken *et al*., 2014; McCracken *et al*., 2017), they do not intrinsically develop a robust mesenchyme. We therefore developed an approach to generate GI mesenchyme from hPSCs that could be incorporated into gastric organoids. Previous work identified a method to direct the differentiation of hPSCs into splanchnic mesenchyme (SM), the source of gastric mesenchyme (Han *et al*., 2020). This method was based on the signaling pathways that drive the normal development of GI mesenchyme and yields a robust population of SM. Briefly, hPSCs were differentiated into lateral plate mesoderm (LPM) with TGFβ inhibition, WNT inhibition, and BMP activation as previously published (Figure 1A) (Han *et al*., 2020; Loh *et al*., 2016). As LPM can give rise to both cardiac and SM, the LPM was treated with retinoic acid (RA) to induce a SM fate, resulting in loss of cardiac markers and an increase in expression of SM markers like *FOXF1* (Figure 1B-C) (Han *et al*., 2020).

**Figure 1.**
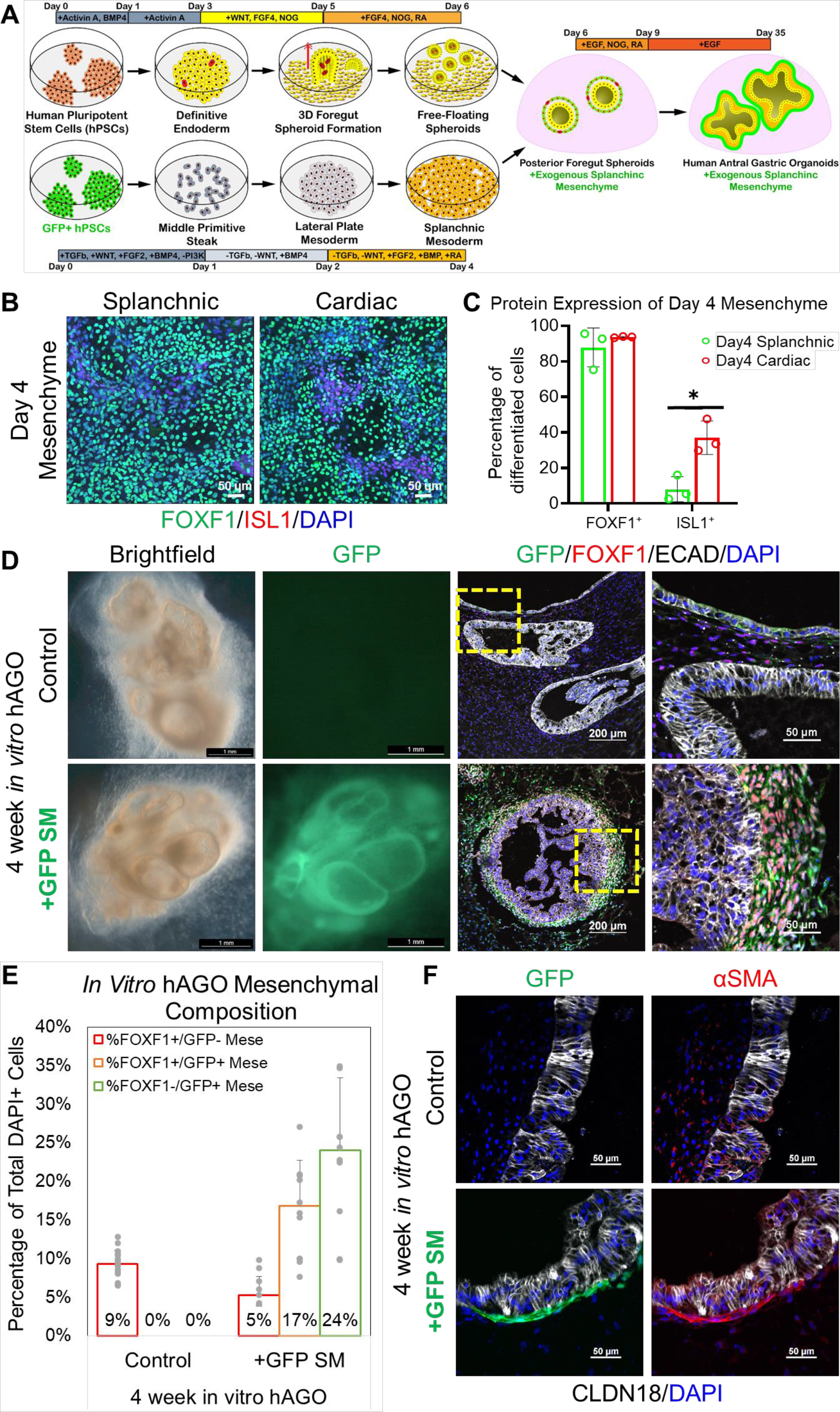
Incorporation of hPSC-derived splanchnic mesenchyme into hAGOs. (**A**) Schematic depicting the method of deriving and incorporating GFP+ splanchnic mesenchyme (SM) into hAGOs. SM was derived from an hPSC line that constitutively expresses GFP. (**B**) Representative immunostaining of day 4 splanchnic (left) and cardiac (right) mesenchymal monolayers costained with FOXF1 (green) and ISL1 (red). (**C**) Quantification of FOXF1+ (left) and ISL1+ (right) cells within day 4 splanchnic (green bar) and cardiac (red bar) mesenchymal monolayers (n=3 fields from one differentiation, *p<0.05, Student’s t-test). (**D**) Brightfield images of hAGOs grown for four weeks *in vitro* with and without recombination with exogenous GFP-labeled SM (green) costained with mesenchymal marker FOXF1 (red). Higher magnification images are shown to the right. (**E**) Quantification of FOXF1+ mesenchymal contribution (n=11-18 fields from at least 3 organoids per condition from one differentiation, same trend seen across at least two individually seeded differentiations, ***p<0.001, Student’s t-test). (**F**) Select images of four week *in vitro* hAGOs with and without recombined SM (green) stained with smooth muscle marker αSMA (red) and gastric epithelial markers CLDN18 (white).

We investigated several approaches to incorporate mesenchyme into developing gastric organoids, including combining mesenchyme and epithelium at different epithelial developmental stages (i.e., at either day 6 or day 9 of the hAGO protocol), testing a single cell aggregation method versus using intact epithelial organoids, and utilizing differently patterned mesenchymal populations, including splanchnic, cardiac, septum transversum, and gastric-esophageal mesenchyme. To monitor this mesenchymal incorporation in real time, we derived the mesenchyme from an hPSC line with constitutive GFP expression. We ultimately found that starting with a single cell suspension of SM that was aggregated with intact posterior foregut spheroids on day 6 of the hAGO protocol (Figure 1A) resulted in optimal mesenchymal incorporation that yielded the most added exogenous mesenchyme while still retaining the small portion of endogenous mesenchyme (Figure S1). To determine this, we initially compared incorporation of varying concentrations of SM and septum transversum (STM) mesenchyme on day 6 of the hAGO protocol. Visual qualitative assessment of brightfield images of 4 week *in vitro* hAGOs recombined with either SM or STM mesenchyme show that recombining SM with hAGO spheroids at a concentration of 50,000 cells/well of approximately 20-30 hAGO spheroids (equating to an approximate 2:1 ratio of SM cells to hAGO epithelial cells) resulted in end timepoint hAGOs +SM that retained an epithelium of visually similar size to hAGO controls (Fig. S1A). We then compared 4 week *in vitro* hAGOs that were recombined with either SM on day 6 of the hAGO protocol or regionally patterned gastric-esophageal mesenchyme (GEM) on day 9 of hAGO protocol (Figure 1D, S1B). After 4 weeks growth *in vitro*, hAGO +SM had a robust and uniform layer of GFP+ mesenchymal cells expressing the early SM marker FOXF1 surrounding the gastric epithelium (Figure 1D), while hAGO +GEM still showed a nonuniform layer of GFP+ mesenchyme (Figure S1B). In gastric organoids +SM, a third to a half of all cells were mesenchyme, representing a 3-5-fold increase over control organoids without added mesenchyme (Figure 1E, S1C). However, in hAGOs +GEM only about a fourth of all cells were mesenchyme (Figure S1C). This was even less for hAGOs +STM (Figure S1C). Overall, mesenchymal populations of SM and CM recombined on day 6 of the hAGO protocol yielded more GFP+ and GFP+/FOXF+ mesenchyme in 4 week *in vitro* hAGOs cultures than populations of GEM and STM that were recombined on day 9 of the hAGO protocol (Figure S1C). Then, between SM and CM populations recombined on day 6, hAGOs +SM retained more endogenous FOXF1+ mesenchyme then hAGOs +CM (Figure S1C). Taken together, we determined that SM recombined with posterior foregut spheroids on day 6 of the hAGO protocol resulted in mesenchymal incorporation that yielded the high populations of both endogenous and exogenous mesenchyme. Finally, in hAGOs +SM we see very rare GFP+ mesenchymal cells that showed the capacity to differentiate *in vitro* into αSMA+ smooth muscle (Figure 1F). We show one image where we see this phenomenon. Otherwise, mesenchymal cells do not differentiate *in vitro* into αSMA+ smooth muscle. This process only occurs after transplantation onto a vascular bed in immunocompromised mice.

Similarly, other aspects of GI organoid growth and morphogenesis are also limited *in vitro* but upon transplantation, intestinal and colonic organoids continue their growth and maturation (Watson et al. 2014). To fully investigate which organoid engineering approach would yield the most characteristic stomach tissue containing glandular units of simple columnar epithelium surrounded by multiple layers of differently oriented smooth muscle layers, all organoids were transplanted into mice and grown for an additional 10 weeks (Table S**1**). Only one organoid of a few mm in size was transplanted into each mouse. Most hAGOs without added mesenchyme did not survive (∼60%) and those that did were small containing only a simple, one-cell layer of thin cuboidal-like epithelium that did not fully differentiate (Figure 2B). In contrast, hAGOs engineered with exogenous mesenchyme had a high survival rate (100%) and grew from a few millimeters to 0.5-1.5 cm in diameter, exhibiting up to 1000x increase in volume in some cases. The mesenchyme differentiated into layers of poorly organized αSMA+ smooth muscle that surround a simple layer of hAGO epithelium (Figure 2B). This data suggests that the incorporation of exogenous mesenchyme promotes engraftment and growth of hAGOs but does not result in normal gastric tissue with glandular units of simple columnar epithelium.

**Figure 2.**
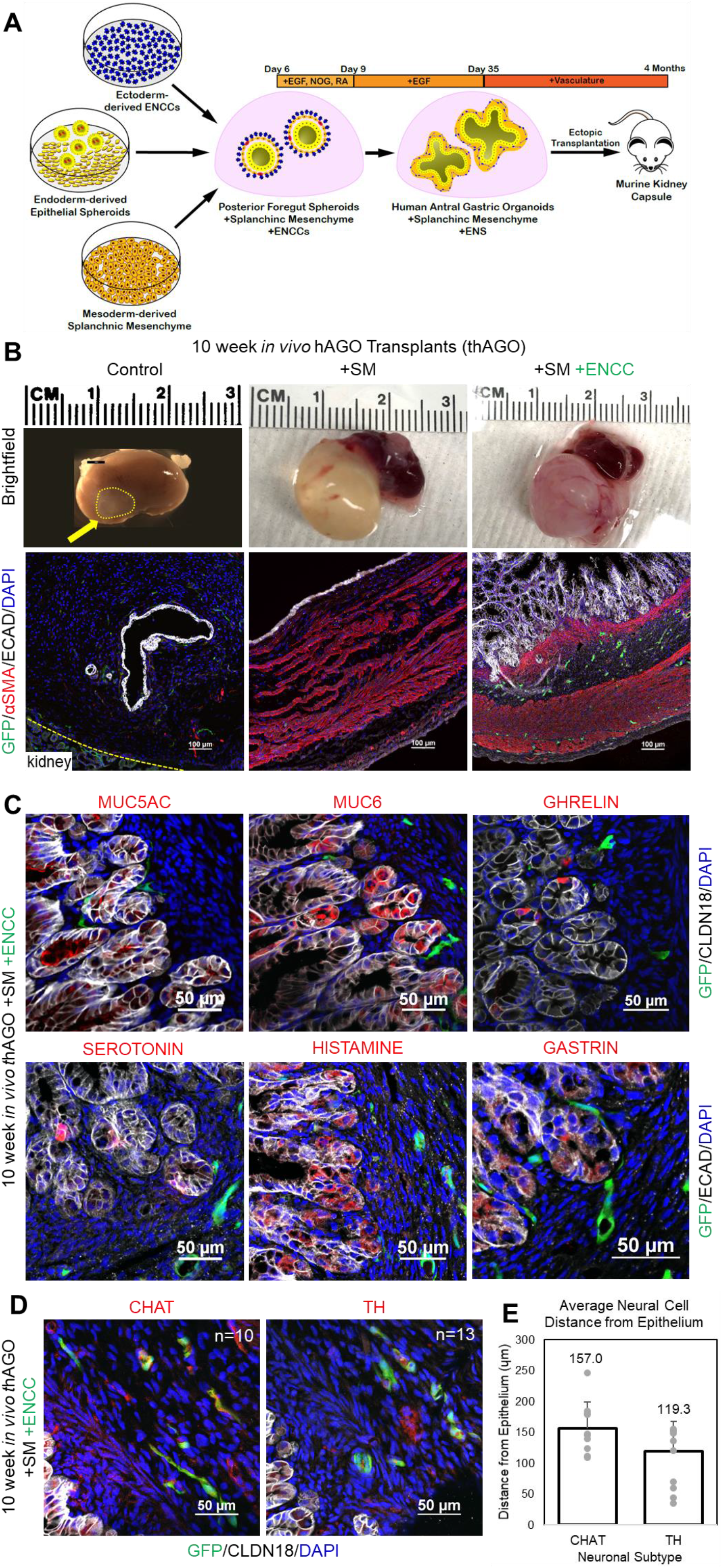
Three germ layer recombinants form human gastric tissue with innervated layers of smooth muscle and glandular epithelium. (**A**) Schematic depicting the generation of three germ layer recombinants using foregut endoderm, SM and ENCCs. (**B**) Morphological comparison between hAGO transplants with and without SM and ENCCs (top) and representative images of 10 week *in vivo* hAGOs stained with αSMA mesenchyme (red) (bottom). ENS is labeled with GFP (green) and counterstained with epithelial marker ECAD (white). (**C**) Marker analysis of gastric epithelial patterning and cell types that develop in three germ layer transplanted hAGOs. MUC5AC (red, top left) and MUC6 (red, top middle) mark surface pit and gland mucous cells, respectfully. Endocrine cells were identified with the hormones ghrelin (red, top right), serotonin (red, bottom left), histamine (red, bottom middle), and gastrin (red, right bottom). GFP labels the recombined ENS (green) and the epithelium is labeled with CLDN18 (white, top) and ECAD (white, bottom). (**D**) Marker analysis of neuronal differentiation in three germ layer transplanted hAGOs. GFP positive ENS (green) is costained with choline acetyltransferase (CHAT, red, left) and tyrosine hydroxylase (TH, red, right). (**E**) Quantification of distance (μm) between epithelium of three germ layer transplanted hAGO and acetyltransferase (CHAT, left) and tyrosine hydroxylase (TH, right) neuronal subtypes (n=10-13 individual cells from two fields from one differentiation). See also Table S1 and Movie S1.

### Engineering three germ layer human gastric tissue

While SM improved the growth of hAGOs, the smooth muscle was poorly organized and there was no evidence of epithelial development into the glandular structures that normally form during human stomach organogenesis. We therefore investigated if incorporation of the ectodermally-derived enteric neural crest cells (ENCCs), in combination with SM, might result in more normal stomach development (Figure 2A). ENCCs migrate into the developing gut tube early in development and form a neuroglial plexus called the enteric nervous system (ENS). There are several published protocols to derived NCCs from hPSCs (Barber, Studer and Fattahi, 2019), but to recapitulate the spatiotemporal dynamics of this developmental process, ENCCs were differentiated as previously described (Workman *et al*., 2017; Bajpai *et al*., 2010) and along with SM were aggregated with hAGO spheroids. By using RFP-labeled ENCCs and GFP-labeled SM we identified conditions that allowed for the incorporation of both germ layers into hAGOs *in vitro* (Figures S2A and S3G). To allow for organoid growth and maturation we transplanted the recombined hAGOs into mice for an additional 10-12 weeks. While both hAGOs +SM and hAGO +SM +ENCC transplants grew to over 1 cm in diameter, only hAGO +SM +ENCC recombinants formed the stereotypic glandular structures found in the human stomach (Figures 2B and 3A-B). In addition, we observed several distinct layers of highly organized smooth muscle that were orientated into distinct layers similar to the organization of the muscularis mucosa, submucosa, and muscularis externa, containing the inner circular and outer longitudinal layers of smooth muscle, of the stomach (Figures 2B, 3A-B, and S2E-F). Our 12wk three germ layer hAGOs are more similar in organization to 38wk human stomach tissue (Figure 3A) when compared to adult human stomach tissue (Figure 3B and S2E-F).” Embedded within the smooth muscle fibers was a network of enteric neurons arranged in characteristic plexi (Figure 2B and Movie S1). The first, more proximal, plexus layer in the 12wk three germ layer hAGO lies in between the more proximal, muscularis mucosa-like muscle layer and the more distal, muscularis externa-like muscle layer, essentially within a submucosal-like space. This plexus layer then is spatially similar to the submucosal neuronal plexus of human stomach tissue. The second, more distal, plexus layer is embedded within the muscularis externa-like muscle layer, mimicking the organization of the myenteric plexus. This organization is highly similar to in vivo 38 week human stomach tissue (Figure 3A and S2E-F).”.

**Figure 3.**
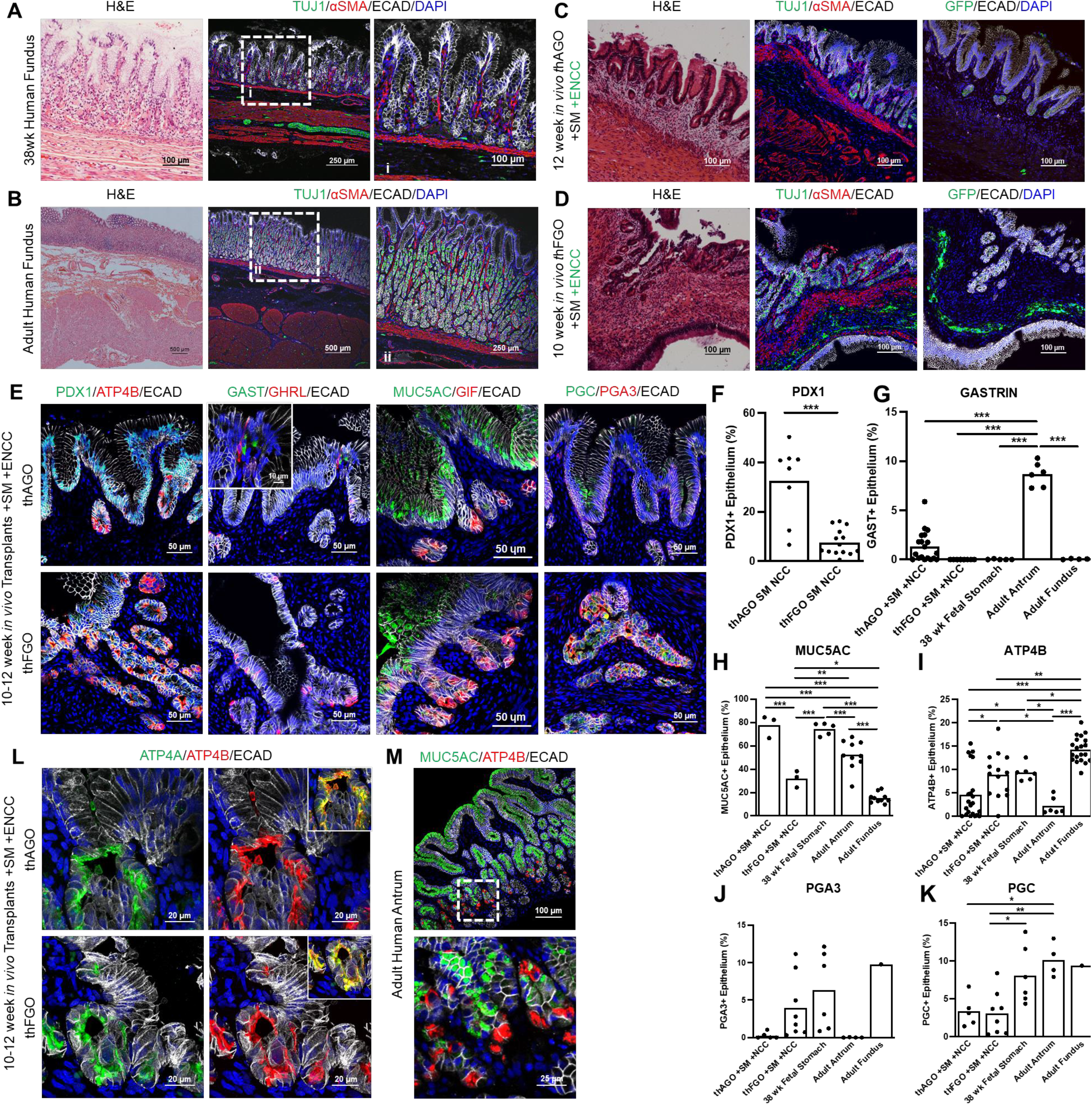
A comparison of engineered antral and fundic organoid tissue with the human stomach. Histological and immunofluorescence analysis of whole thickness gastric tissue from 38wk (**A**) and adult (**B**) human stomach taken from the distal fundus and three germ layer transplanted (**C**) hAGO and (**D**) hFGO. The three germ layers are labeled with neuronal marker TUJ1 or GFP (green), smooth muscle αSMA (red), and epithelial marker ECAD (white). (**E**) Representative images of gastric epithelial patterning and cell differentiation of three germ layer transplanted hAGOs (top) and hFGOs (bottom). PDX1 (green, left), endocrine cells expressing gastrin (GAST, green, middle), and the surface mucin (MUC5AC, green, middle) are (**E**) qualitatively and (**F-H**) qualitatively enriched in hAGOs and human antral biopsies. Parietal cells (ATP4B, red, left; and GIF, red, middle and chief cells expressing fundic-specific pepsinogen A3 (PGA3, red, right) are (**E**) qualitatively and (**I-J**) quantitatively enriched in the hFGOs and human fundic biopsies. Endocrine cells expressing ghrelin (GHRL, red, middle) and chief cell marker pepsinogen C (PGC, green, right) are observed at relatively similar levels in both thAGOs and thFGOs (**E**,**K**). (**L**) Representative images of parietal cells in three germ layer transplanted hAGOs (top) and hFGOs (bottom) with colocalized apical expression of ATP4A (green) and ATP4B (red) H+/K+ ATPase subunits. (**E, L, M**) Epithelium is labeled with ECAD (white). Significance denoted as *p<0.05, **p<0.01, and ***p<0.001 determined by (**F**) Student t-test or (**G-K**) one-way ANOVA with Tukey’s Multiple Comparison (n=3-20 fields from 5 thAGO TXPs from two differentiations, 8-14 fields from 2 thFGO TXPs from one differentiation, 5-6 fields from fetal stomach from 1 patient, 1-20 fields from adult fundus from 3 patients, and 4-10 fields from adult antrum from 2 patients). (**M**) Representative human adult antral biopsy labeled with MUC5AC (green) and ATP4B (red). See also Figures S2 and S3.

The epithelium of hAGO +SM +ENCC transplants expressed the gastric epithelial marker CLDN18 and lacked the intestinal epithelial marker CDH17, confirming the gastric identity of the organoids (Figure 2C, data not shown). hAGO glands contained all of the expected cell types normally found in the antrum of the stomach including surface mucous cells (MUC5AC), gland mucous cells (MUC6), and endocrine cells expressing ghrelin, serotonin, histamine, and gastrin (Figure 2C and 3E). The neurons (GFP+) formed a network of fibers resembling a plexus that was embedded within the layers of smooth muscle. We also observed GFP+ choline acetyltransferase+ (ChAT+) and dopaminergic (TH+) neurons approximately 120-160μm away from the glandular epithelium and in close proximity to the endocrine cells such as ghrelin and gastrin (Figure 2C-E). This association *in vivo* is important as neurotransmitters control secretion of a variety of stomach hormones including ghrelin and gastrin (Breit *et al*., 2018).

### Generating fundic tissues containing three germ layers

One of the most prominent domains of the human stomach is the corpus, which contains fundic (oxyntic) glands with acid producing parietal cells and digestive enzyme secreting chief cells. The glands are also in close proximity to enteric neurons that, along with gastric endocrine cells, help to regulate acid production. We investigated if the three germ layer recombinant approach could also be used to engineer human fundic tissue with the above properties. We generated early stage hFGOs as previously described (McCracken *et al*., 2017) and recombined them with SM and GFP-labeled ENCCs. After four weeks we observed both SM and ENCCs incorporated into hFGOs (Figure S2H), similar to hAGOs (Figures S2A-D), and confirmed fundic identity by the presence of ATP4B+ parietal cells and absence of PDX1 (Figure S2H and data not shown).

We next investigated if, like hAGOs (Figure S2C), incorporation of SM and ENCCs also promoted growth, morphogenesis, and maturation of hFGOs engrafted under the murine kidney capsule (Figure S2G). A comparison of three germ layer hAGOs and hFGOs grown *in vivo* for 10-12 weeks showed that they both grew up to a centimeter in size (Figure S2C and S2G) with a similar histological architecture to that of 38 week (Figure 3A) and adult (Figure 3B) human fundic tissues, with glandular epithelium surrounded by multiple layers of innervated smooth muscle (Figure 3A-D. In general, the extent of glandular morphogenesis of transplanted hFGOs was less than that of hAGOs (Figure 3C-E). Both hAGOs and hFGOs maintained their regional identity after transplantation, and moreover, the proportions of cell types that normally distinguish the human corpus/fundic from the antrum also distinguished hAGOs from hFGOs. Specifically, hAGOs expressed higher levels of PDX1, antral-specific gastrin-expressing endocrine cells, and MUC5AC+ surface mucus cells compared to hFGOs (Figure 3E=H). Conversely, hFGOs contained more ATP4B+/GIF+ parietal cells than the hAGOs, and hFGOs had fundic-specific PGA3+ chief cells, which were absent in hAGOs (Figure 3E, 3I-J). Ghrelin-expressing endocrine cells and PGC+ chief cells were observed in both regions of the stomach (Figures 3E, 3K). We previously demonstrated that differentiation of parietal cells *in vitro* required BMP signaling and MEK inhibition (Figure S2H) (McCracken *et al*., 2017), however transplanted hFGOs required no additional factors for robust parietal cell differentiation (Figures 3E, 3I, and 3L-M) demonstrating that the signaling processes that control gastric cell type specification occur normally in engineered tissue. As observed in human stomach biopsies, engineered antral tissue does contain parietal cells, but at lower numbers than are found in fundic glands (Figures 3E, 3I, and 3L-M) (Choi *et al*., 2014). Furthermore, parietal cells in hFGOs *in vitro* only expressed the ATP4B+ subunit of the H+/K+ ATPase and the cellular localization is primarily cytoplasmic (Figure S2H) while hFGOs matured *in vivo* expressed much higher levels of both the ATP4A and ATP4B subunits which colocalize on the apical membrane (Figure 3L) suggesting that these parietal cells are more mature than their *in vitro* counterparts. Together these data confirm that engineering mesenchyme and ENS cells into hAGOs and hFGOs results in the formation of gastric tissues that begin to resemble human stomach tissue.

### Antral three germ layer organoids exhibit functional muscle contraction

The stomach plays an essential role in the mechanical breakdown of food and in emptying it into the duodenum. This gastric motility involves the ENS, which functionally controls smooth muscle contractions. To investigate if the ENS and smooth muscle in the three germ layer hAGOs formed a functional neuromuscular unit, we isolated tissue strips from transplanted hAGOs and placed them in an organ bath chamber system to monitor contractility. After an equilibration period, spontaneous contractile oscillations were observed from tissues derived from 1 hAGO +SM and 3 separate hAGO +SM +ENCC transplants (Figure 4A). The presence of phasic contractions indicated that intramuscular interstitial cells of Cajal (ICCs) were present within both the two and three germ layer organoids. However, the contractile activity exhibited in hAGO +SM had more irregularities than observed in hAGO +SM +ENCC. This was further supported by the presence of mesenchymal clusters expressing KIT Proto-Oncogene, Receptor Tyrosine Kinase (c-KIT), a marker of ICCs, that were in close association with TUJ1+ neuroglia cells (Figure 4B), indicating cooperative coordination of the contractions in hAGO +SM +ENCC improved their regularity (Ward and Sanders, 2006; Iino and Horiguchi, 2006). These mesenchymal clusters, approximately 7% of total cells present (Figure 4C), arranged within the muscularis externa-like muscle layers of the three germ layer hAGO transplants as is stereotypical in the human stomach. Smooth muscle tone was then interrogated with a dose response to bethanechol, a muscarinic receptor agonist that directly stimulates smooth muscle contractions (Figure 4D). The contractility increased in response to bethanechol in a dose-dependent manner, demonstrating the presence of functional smooth muscle in both hAGO +SM and hAGO +SM +ENCC. Moreover, we were able to reverse the contractions and induce muscle relaxation with addition of scopolamine, a muscarinic antagonist, in both groups (Figure 4E). Taken together, these data were indicative of functional muscle tissue in the *in vivo* engrafted hAGOs.

**Figure 4.**
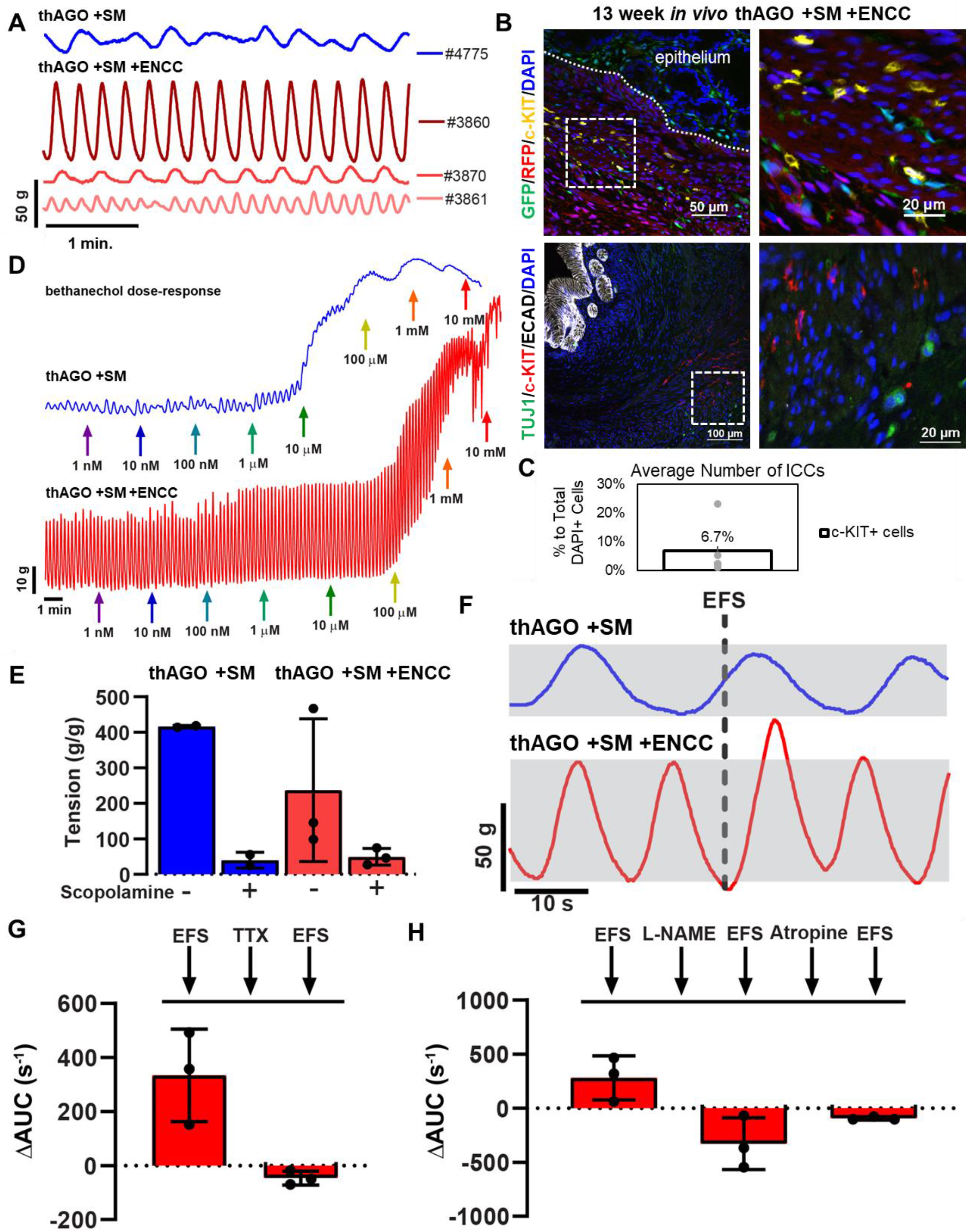
Antral three germ layer organoids have a functional ENS that regulates gastric tissue contractions. (**A**) Isometric force contractions in tissues isolated from one individual transplanted hAGO +SM (blue) and three individual transplanted hAGO +SM +ECC (red) after an equilibrium period. Contractile activity was triggered using electrical field stimulation (EFS). (**B**) Neuronal (GFP, green, top; and TUJ1, green, bottom) and interstitial cells of Cajal (ICC) (c-KIT, yellow, top; and red, bottom) stained in 13 week *in vivo* hAGO +SM +ENCC grafts. (**C**) Quantification of the average number of ICCs in *in vivo* hAGO +SM +ENCC grafts (n=7 fields from 3 organoids from 2 differentiations). (**D**) Activation of muscarinic receptors induced contractions in tissues isolated from a transplanted hAGO +SM (blue) and hAGO +SM +ECC (red). Increasing doses of bethanechol were added to the tissues at times indicated by the colored arrows. (**E**) Inhibition of the muscarinic receptor with scopolamine induced muscle relaxation. Calculated maximal and minimal tissue tension of tissues from hAGO +SM (blue) and hAGO +SM +ECC (red). (**F**) Representative tracings of contractile activity in response to electrical field stimulation (EFS) in transplanted hAGO +SM (blue) and hAGO +SM +ENCC (red). Dashed line indicates timing of EFS application and gray rectangles highlight pre-EFS contractile amplitude. (**G**) Inhibition of ENS activation with the neurotoxin tetrodotoxin (TTX) abrogates EFS-mediated contractions. Change in area under the curve following a control EFS stimulation measured for one minute after stimulation, followed by TTX treatment, and a final EFS stimulation in hAGO +SM +ENCC. (**H**) Functionally testing the role of nitrergic and cholinergic neuronal activity in smooth muscle contractions. Change in area under the curve induced by EFS stimulation and following treatment with the nitrergic inhibitor L-NAME and the cholinergic inhibitor Atropine. All data was normalized to tissue mass; n=1 hAGO +SM; n=3 hAGO +SM +ENCC from two differentiations.

We next investigated if the ENS that we engineered into hAGOs was functionally capable of controlling gastric tissue contractions. Electrical field stimulation (EFS) of tissues is an experimental means to trigger neuronal firing and subsequent smooth muscle contraction. EFS pulses were administered to two and three germ layer hAGO muscle strips and only resulted in an increase in contractile activity in hAGO +SM +ENCC, indicating that the ENS was regulating smooth muscle (Figure 4F). To show that there was a functional connection between the ENS and smooth muscle, we inhibited ENS activity with the neurotoxin tetrodotoxin (TTX), which abolished the ability of EFS to stimulate contractile activity (Figure 4G). Lastly, we investigated the involvement of nitrergic and cholinergic neuronal activity in regulating smooth muscle contractions. We inhibited nitric oxide synthetase (nNOS)-expressing neurons with NG-nitro-l-arginine methyl ester (L-NAME), a nitric oxide synthesis inhibitor, and inhibited cholinergic neurons using atropine, an acetylcholine (Ach) receptor antagonist. Contractile activity was measured following control stimulation and stimulation after compound exposure and was expressed as the change in the area under the curve (AUC) immediately before and after each EFS stimulation (Figure 4H). These data provide insight into the proportions of nitrergic and cholinergic neuronal activity compared to the total ENS activity (control EFS) and show that gastric tissue contractions involved both nitrergic and cholinergic neuronal activities.

### Three germ layer esophageal organoids

To test whether our approach of combining tissue from three germ layers was broadly applicable to engineering other organs, we attempted to incorporate SM and ENCCs into developing human esophageal organoids (HEOs) (Trisno et al., 2018). Like hAGOs and hFGOs, we started with HEOs that are largely epithelial and added GFP-labeled SM (Figure S3A-B). After four weeks *in vitro* HEOs +SM had a robust layer of GFP+ mesenchyme surrounding the epithelium (Figure S3A) with a high percent of these co-expressing FOXF1 or the more differentiated marker vimentin (VIM) (Figure S3A). Quantification showed that control HEOs only contain ∼1% of endogenous mesenchyme while HEOs +SM contain ∼25% mesenchymal cells (Figure S3B). Interestingly the addition of exogenous, GFP+ mesenchyme facilitated the expansion of endogenous FOXF1+/GFP-mesenchyme in the cultures, suggesting cell-cell interactions promote the growth and development of both the organoid epithelium and mesenchyme.

We next incorporated ENCCs into HEOs with or without SM (Figure S3C-H). After 1 month of *in vitro* culture, the ENCCs in HEOs without SM had differentiated into TUJ1/MAP2/NESTINM+ enteric neurons that aggregated tightly around the epithelium and did not organize into a neuronal plexus (Figure S3D-E). In contrast when both ENCCs and splanchnic mesenchyme were recombined with HEO epithelium we did observed robust co-development of TUJ1+ neuronal plexus associated within FOXF1+ mesenchymal layer (Figure S3F-I). Overall, these finding show that different human GI organ tissues can be engineered by combining progenitors from all three germ layers and emphasize the importance of reciprocal cell-cell communication between the epithelial, mesenchymal, and ENCCs for proper assembly and function of embryonic organs.

### ENCCs differentiation into ENS neuroglial cell types does not require the addition of exogenous mesenchyme

One of the most powerful aspects of this system is the ability to study interactions between cell types of different germ layers that drive normal tissue formation. For example, our findings suggested that the presence of ENCCs was important for the development of both the smooth muscle and the gastric epithelium. Without ENCCs, mesenchyme formed a small layer of disorganized smooth muscle and the gastric epithelium failed to undergo glandular morphogenesis. We decided to take advantage of the fact that we can add or remove germ layers at will to interrogate how ENCCs impact the development of the other germ layers. We first independently differentiated hPSCs into migrating vagal-like ENCCs (Figure S4A) that after two weeks *in vitro* expressed key ENCCs markers, including SOX10, AP2A, and p75 (Figure S4C), and upregulated key neural crest specifier genes SOX9, SOX10, and SNAIL2 (Figure S4D). We then recombined ENCCs with hAGOs at two different timepoints, day 6 and day 9 of gastric organoid development, and determined their ability to form ENS cell types without exogenous mesenchyme (Figure 5A). The rationale for recombining ENCCs at day 9 was to avoid exposing ENCCs to retinoic acid (RA) and noggin (NOG) that are in the hAGO cultures between days 6-9 as it was previously shown that addition of RA to ENCCs in monolayer culture posteriorized their axial identity as indicated by upregulation of regional Hox genes HOXB3, HOXB5, and HOXB7 (Fig S4E) (Simoes-Costa and Bronner, 2015; Workman *et al*., 2017). Surprisingly at either time point, ENCCs incorporated well into hAGOs and formed a 3D network of TUJ1+ neurons and S100b+ glial cells adjacent to gastric epithelium (Figure S4B, F-H and Movie S2). ENCCs differentiated into a diverse array of neuroglial subtypes, including inhibitory (nNOS), interneurons (Synaptophysin), dopaminergic (TH), sensory (Calbindin) neurons, and glial cells (GFAP) (Figure S4I and Table S2). GFAP cells represent approximately 2% of the total cells present within 4 week *in vitro* hAGOs +ENCC (Figure S4J). ENCCs did not alter gross hAGOs growth or morphology after four weeks of development *in vitro* (Figure S4B). However, ENS development was abnormal. Neurons were found immediately adjacent to the gastric epithelium and were disorganized as compared to mouse E13.5 embryonic stomach (Figure S4K-L). There were, however, a comparable number of nNOS+ inhibitory neurons present in hAGOs +ENCC compared to mouse E13.5 stomach (Figure S4M-N). These data show that ENCCs incorporated into hAGOs differentiated into neuroglial subtypes without the addition of exogenous mesenchyme, but that proper spatial orientation and ENS plexus development likely requires a robust population of mesenchyme.

**Figure 5.**
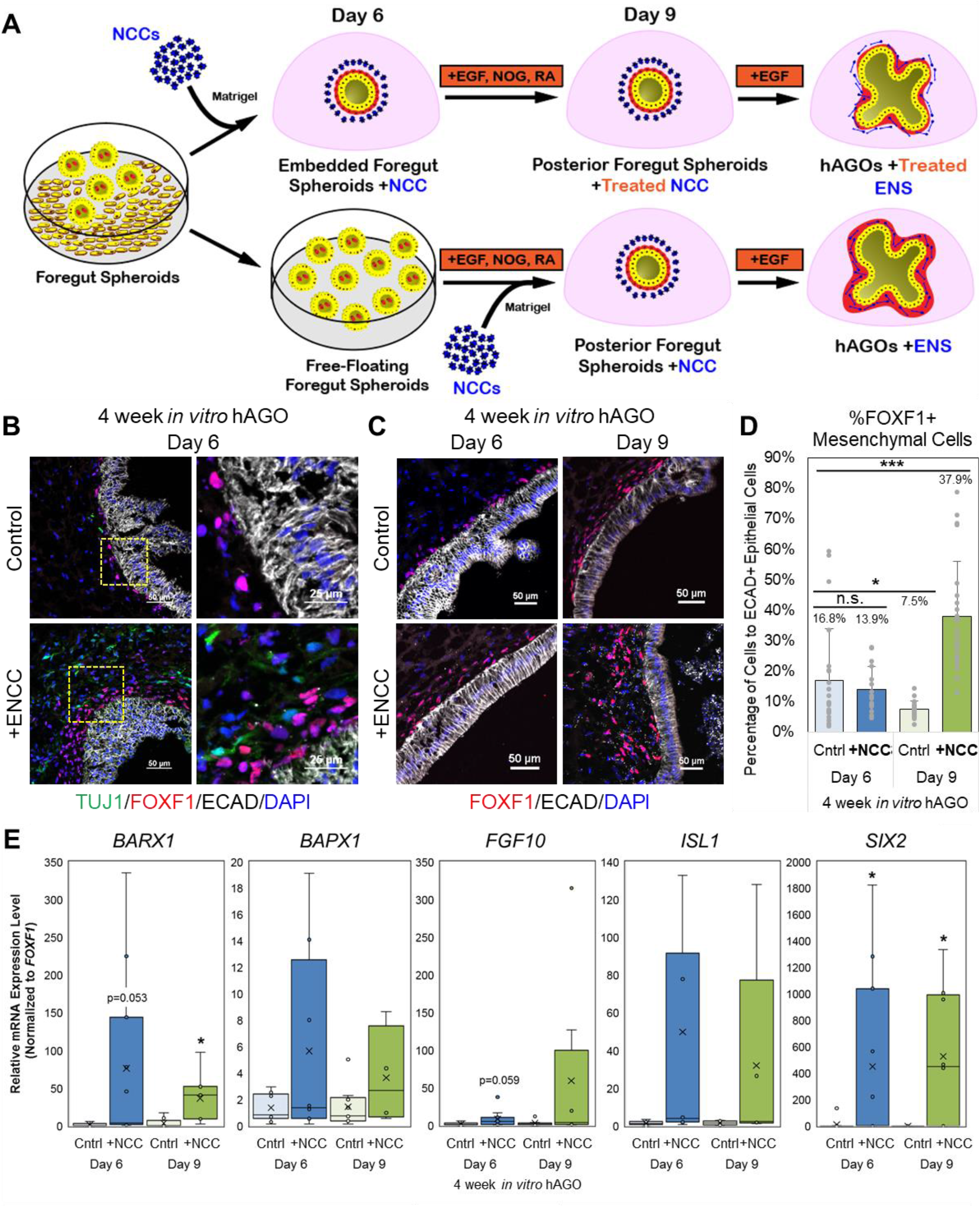
ENS cells promote *in vitro* growth and patterning of gastric mesenchyme. (**A**) Schematic illustrating the method of recombining hAGOs with ENCCs at day 6 and day 9 of hAGO protocol. (**B**) Representative images of four week *in vitro* hAGOs with (bottom) and without (top) ENS recombined on day 6 of hAGO protocol stained with TUJ1 neurons (green) and FOXF1 mesenchyme (red) and epithelial ECAD (white). Higher magnification images are shown to the right. (**C**) Representative images of four week *in vitro* hAGOs with (bottom) and without (top) ENS recombined on either day 6 (left) or day 9 (right) of hAGO protocol, demonstrate an increase in FOXF1+ mesenchyme (red). (**D**) Quantification of FOXF1+ mesenchymal contribution (n=16-24 fields from at least 3 organoids from one differentiation, same trend seen across at least two individually seeded differentiations, *p<0.05, ***p<0.001, Student’s t-test). (**E**) Relative expression of key gastric mesenchymal genes (*BARX1, BAPX1, FGF10, ISL1, SIX2*) in hAGOs +ENCC when compared to hAGOs -ENS. (n=4-12 wells, with a minimum of 3 organoids per well, from 5 individual differentiations, *p<0.05, Student’s t-test). See also Figure S3, Movie S2, and Table S1.

### ENS cells promote the growth and gastric identity of mesenchyme

Previous studies in developing chicken embryos suggest that ENCCs are involved in gastric mesenchyme development (Faure *et al*., 2015). We therefore analyzed the impact of added ENCCs on the development of the small amount of endogenous mesenchyme present in hAGOs. Addition of ENCCs at day 6 of hAGO development had little effect on the number of FOXF1+ mesenchyme cells; in contrast, addition of ENCCs at day 9 resulted in 2-4 times more FOXF1+ mesenchyme surrounding the epithelium (Figure 5A-D). Addition of ENCCs at day 6 or day 9 also correlated with increased levels of gastric mesenchymal genes *BARX1, BAPX1, FGF10, ISL1*, and *SIX2* (Figure 5E) (Faure *et al*., 2013). This suggested that the enteric neurons not only encourage the growth of mesenchyme *in vitro*, but also support its proper regional patterning into gastric-specific mesenchyme. It is interesting that addition of ENCCs at day 9 promotes the expansion of mesenchyme whereas addition at day 6 does not. The main difference is that ENCCs recombined with hAGOs at day 6 are exposed to the BMP inhibitor NOG and RA from day 6-9 as part of the normal hAGO protocol. The impact of this treatment on ENCCs will be discussed below.

### ENCC cells support the growth and morphogenesis of organoid epithelium *in vivo*

We described above (Figure 2) that addition of exogenous mesenchyme alone was not sufficient to promote growth and morphogenesis of organoid epithelium. However, we did not investigate how the addition of ENCCs alone without exogenous mesenchyme might impact epithelial development. Therefore, hAGOs with ENCCs recombined at day 6 and 9 were transplanted into mice and grown for 6-15 weeks at which time they were scored for graft survival, overall growth, and epithelial morphogenesis (Figure S5A-B). The presence of an ENS, even in the absence of exogenous mesenchyme, improved both number and epithelial growth of hAGO +ENCC grafts (Figures 6A-B and S5B-D). In most cases the epithelium of the grafts was a simple gastric epithelium with gastric hormonal cells, such as gastrin, ghrelin, somatostatin, and serotonin, as well as surface mucous cells marked by MUC5AC (Figure S5E). However, in 5/21 hAGO +ENCC grafts we observed pronounced glandular epithelial morphogenesis as compared to 0/19 hAGO -ENCC grafts (Figure 6A-B). A time course analysis of grafts 4, 10, and 14 weeks following transplantation showed rare examples of differentiated smooth muscle (Figure S5F) and neuroglial cells expressing TUJ1, S100b, peripherin, nNOS, and GFAP (Figure S6A-B). GFAP cells represent approximately 1% of the total cells present within 14 week *in vivo* hAGOs +ENCC (Figure S6C). These neurons are arranged near the epithelium (Figure S6D and Movie S3) and capable of effluxing calcium as measured using a GCaMP reporter as previously shown (Figure S6E-F and Movie S4) (Workman *et al*., 2017; Chen *et al*., 2013). Together, these data indicate that ENCCs promote survival and engraftment of hAGOs and the development of glandular tissue in subset of grafts. However, without a sufficient amount of mesenchyme, addition of ENCCs alone will not result in the development of normal gastric tissue.

**Figure 6.**
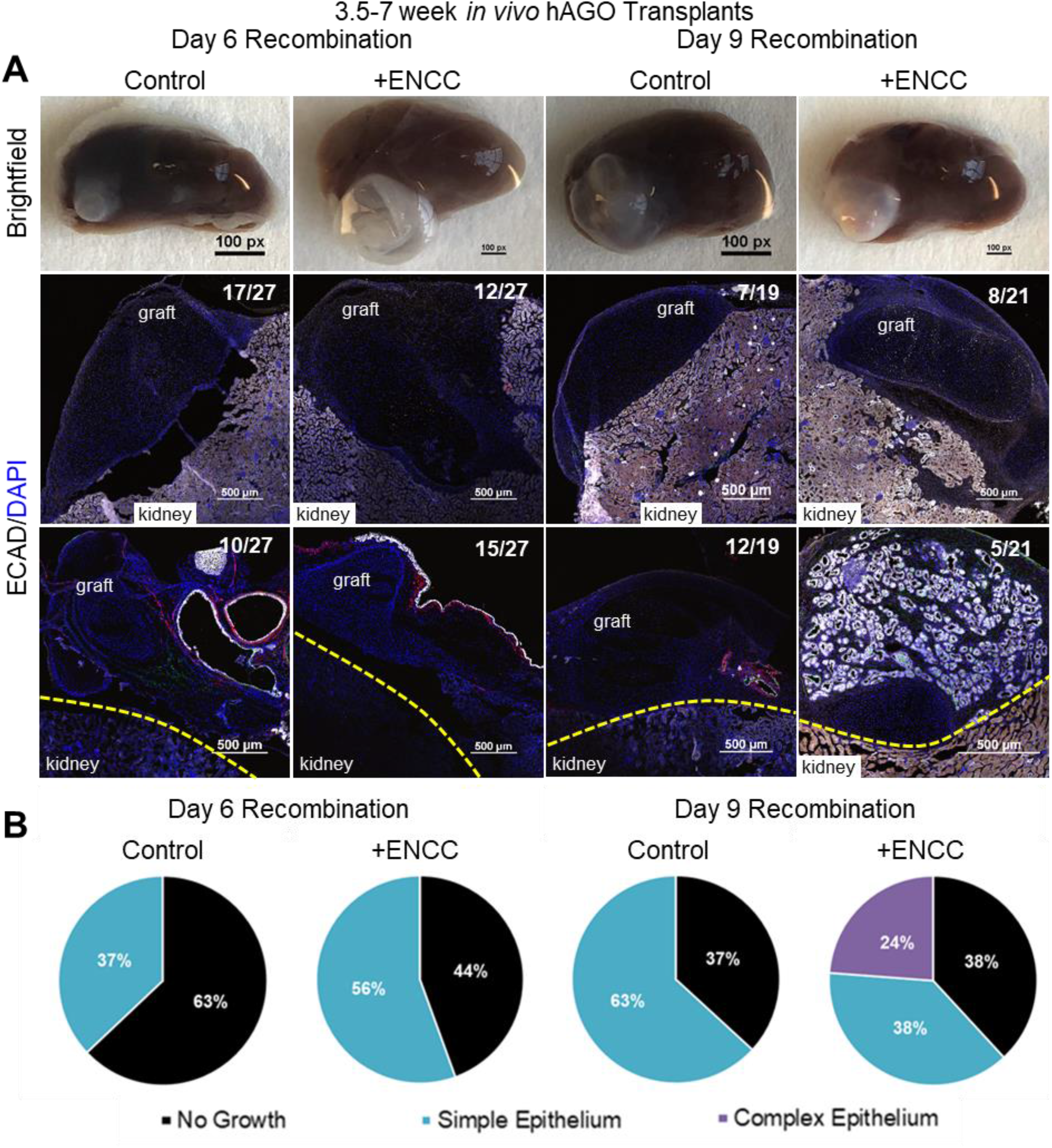
ENCCs promote hAGO engraftment and epithelial growth. (**A**) Representative low magnification images of gross organoids of ECAD+ epithelium (white) from transplanted hAGOs with and without ENS following recombination at day 6 or day 9. (**B**) Quantification of organoid engraftment and epithelial growth from transplanted hAGOs with and without ENCCs. 24% (5/21) of hAGO +ENCC recombined at day 9 had complex glandular epithelium. See also Figures S5 and S6 and Movies S3 and S4.

### The epithelium of hAGOs +ENCCs is morphologically and molecularly similar to Brunner’s Glands

A number of hAGO +ENCC grafts displayed a complex glandular epithelial morphology (Figures 6A-B and S7C-D), expressed PDX1 and GATA4 indicative of gastrointestinal regional identity, and had hormone-expressing cells such as serotonin, ghrelin, histamine, and somatostatin (Figure 7A, data not shown). However, they did not express key gastric-specific epithelial markers CLDN18 or SOX2 (Figure 7A-B) or have characteristic gastric cell types MUC5AC-expressing mucous cells (Figure 7D). The glandular epithelium was also negative for intestinal epithelial markers CDX2 and CDH17 (Figure 7A-B). Lastly, we confirmed that these were human tissue and not a contaminant mouse tissues from the host (data not shown). Given that the glandular epithelium of the grafts was neither gastric nor intestinal, we explored the possibility that these were Brunner’s glands. Brunner’s glands are glandular structures found within the submucosa of the proximal part of the duodenum, near the pyloric junction. They serve to secrete sodium bicarbonate to neutralize any escaping gastric acids. Given the lack of definitive markers for human Brunner’s glands, we established a combinatorial marker profile for Brunner’s glands using patient biopsies (Figure S7A-B) and published reports (Figure 7C). Human Brunner’s glands are negative for gastric markers CLDN18, SOX2, and MUC5AC and intestinal markers CDH17, MUC2 and have low levels of CDX2 compared to adjacent duodenal epithelium (Figure S7). Human Brunner’s glands are positive for glucagon-like peptide-1 receptor (GLP-1R) and MUC6 and co-expression of these markers occurs only in Brunner’s glands. The combinatorial expression profile of 9 different markers supports the conclusion that the glandular epithelium of hAGO +ENCC grafts is most similar to Brunner’s glands (Figures 7C and S7A-B) (Tan *et al*., 2020; Wang *et al*., 2015; Balbinot *et al*., 2017).

**Figure 7.**
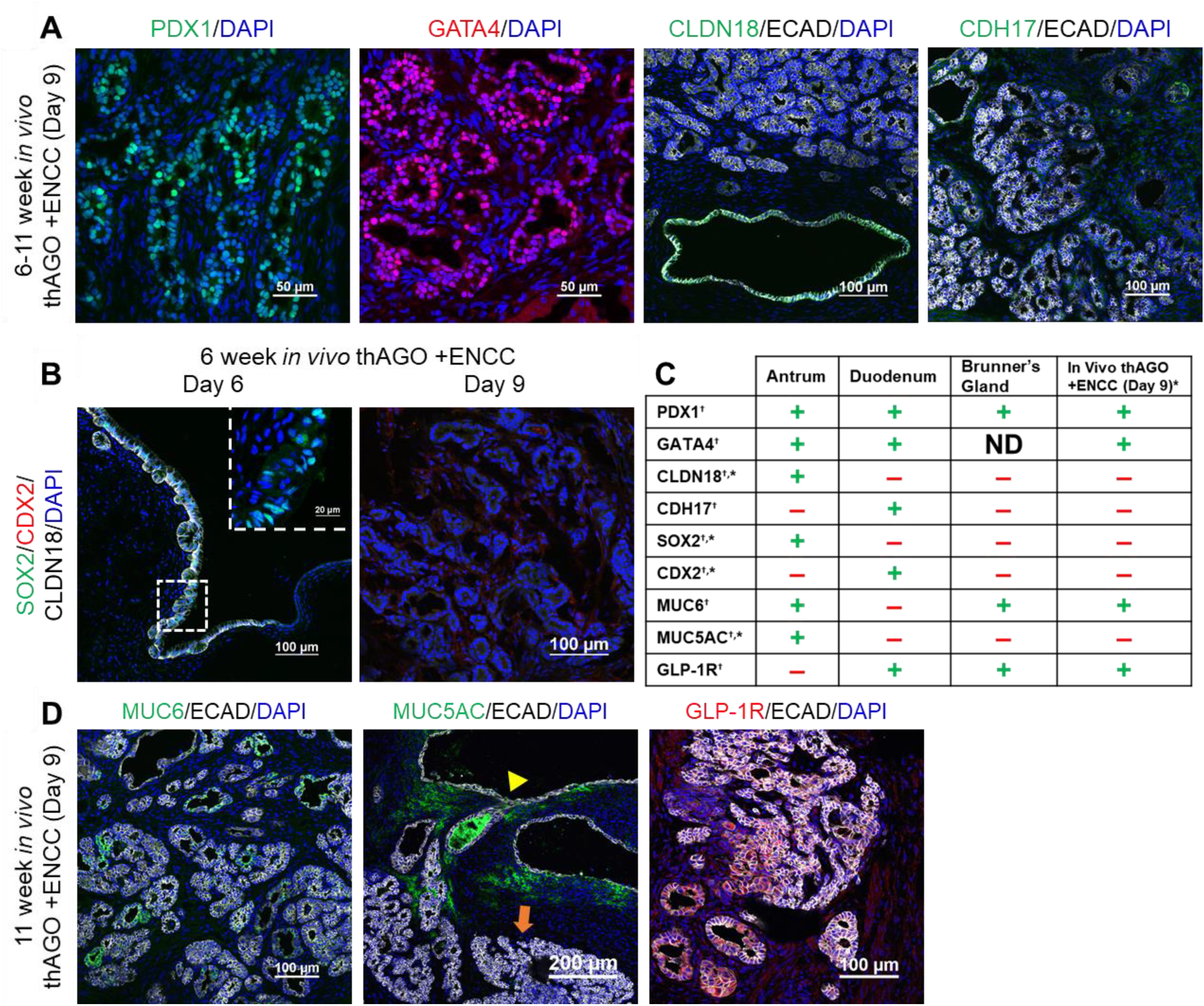
Identification of hAGO +ENCC glandular epithelium as Brunner’s Glands. (**A**) Glandular epithelium expressed the pan gastrointestinal markers PDX1 (green) and GATA4 (red) but did not express gastric epithelial marker CLDN18 or intestinal epithelial marker CDH17 in transplanted hAGOs recombined with ENCCs on day 9. (**B-D**) Marker analysis of organoid epithelium at different time points following transplantation. (**B**) After 6 weeks growth *in vivo* hAGOs with ENCCs recombined at day 6 had a simple epithelium expressing the gastric markers SOX2 (green, inset) and CLDN18 (white) but not the intestinal marker CDX2 (red). The glandular epithelium of from day 9 recombinants did not express these gastric or intestinal markers. (**C**) Comparison of antral and duodenal markers to known markers of Brunner’s glands and how these align with observed protein expression profile of complex epithelial growths from hAGOs +ENCC day 9 recombined grafts. ^†^determined from previously published data; *determined experimentally on human tissue samples of Brunner’s Glands (see Figure S7B) (**D**) At 11 week post-transplant, the glandular epithelium in organoids did express MUC6 (green, left) and GLP-1R (green, middle), similar to human Brunner’s glands. The simple epithelium (yellow arrowhead) expressed MUC5AC (green, right) while complex epithelium (orange arrow) the glandular epithelium did not. See also Figures S7 and S8.

This shift in hAGO epithelium from gastric identify to the more posterior Brunner’s gland identity suggest that the added ENCCs were driving this more posterior fate, suggesting that ENCCs, in the absence of mesodermal contribution, may produce a factor(s) that posteriorize gastric epithelium. One candidate pathway was BMP signaling, which is known to promote posterior fate in the gastrointestinal junction (Smith and Tabin, 1999; Smith *et al*., 2000; Faure *et al*., 2002; Tiso *et al*., 2002; De Santa Barbara *et al*., 2005; Theodosiou and Tabin, 2005; Davenport *et al*., 2016; Stevens *et al*., 2017). Analysis of ENCCs show high levels of expression of both *BMP4* and *BMP7* (Figure S8A). To functionally investigate if BMP activity might mediate the ability of ENCCs to promote a Brunner’s gland fate, ENCCs were recombined with hAGO at day 6 and then organoids were cultured with the BMP inhibitor NOG from day 6-9, along with RA, which is a component of the normal hAGO protocol. None of the grafts (0/27) had Brunner’s gland epithelium following 3 days of noggin treatment as compared to hAGOs +ENCCs grafts that were not treated with NOG, where the 5/21 grafts contained Brunner’s gland epithelium. To investigate if NOG-treated ENCCs might have reduced posteriorizing activity we treated ENCC cultures with NOG and found significantly reduced levels of both *BMP4* and *7* (Figure S7B). We conclude that posteriorizing factors like *BMP4* and *7* are produced by ENCCs and that in the presence of BMP inhibitors, ENCC lose their ability to posteriorize gastric epithelium.

## DISCUSSION

By understanding and applying the key signaling pathways known to regulate the development of different cell types, many protocols have now been published to direct the differentiation of hPSCs into germ layer-specific fates, including endoderm-derived epithelial cells, mesoderm-derived mesenchymal cells, and ectoderm-derived neural cells. This research extended those principles to also apply the known spatiotemporal events for GI organ assembly to tissue engineering, starting from separately derived germ layer derivatives. From studies of embryo development, we know that the GI tract is assembled in a step wise manner. First a two-dimensional sheet of endoderm forms a gut tube that becomes encapsulated by splanchnic mesenchyme that is then invaded by ENCCs that have migrated from the neural tube. This all happens within a few days during embryonic development. We mimic this by instructing a 2D sheet of endoderm to undergo morphogenesis forming gut tube-like spheroids, and then physically surrounding them with a cellular mixture of SM and ENCCs. To our knowledge, this co-culture of three unique cell types that grew together to form three germ layer organoids is the best hPSCs-derived approximation of bona fide human stomach tissues. The basic concept of assembling organoids from separately-derived germ layer progenitors was also applied to both fundus and esophagus, suggesting that this technology could be broadly applied to tissue engineer other organs, like lung, liver, and bladder.

Congenital diseases in humans often affects several organs and can be due to impacts on multiple germ layers. These three germ layer organoids represent new model systems to study both the effects of patient-specific mutations on multiple organs and how gene mutations impact individual germ layers, similar to a cell specific Cre approaches in mice. This approach has been used to study the impact of patient mutations on human PSC-derived ENCCs (Workman *et al*., 2017), on PSC-derived epithelial cell types (Zhang *et al*., 2019), and could be used to study mutations that largely effect mesenchyme (Gilbert *et al*., 2020). Analyses of organoids could be used to identify previously unappreciated patient pathologies that could inform improved clinical care. However, to effectively understand congenital disorders, we need to better understand the processes of epithelial-mesenchyme-ENS communication during normal organ development. The tractability of this system seems ideally suited to interrogate such signaling pathways mediating this crosstalk. Our data suggests that ENCCs impact the growth and patterning of endogenous mesenchyme. Moreover, without addition of exogenous splanchnic mesenchyme, ENCCs re-pattern gastric epithelium to a more posterior identity similar to Brunner’s glands. However, when ENCCs are added simultaneously with a robust population of mesenchyme, together these germ layers maintain gastric patterning and promote gastric gland morphogenesis.

The ability to manipulate signaling pathways at will and in a germ layer-specific manner *in vitro* is a powerful way to dissect the molecular basis of organ development. For example, it is known that the WNT and BMP signaling pathways control the anterior-posterior and dorso-ventral patterning of the developing GI tract in model organisms (Smith and Tabin, 1999; Smith *et al*., 2000; Faure *et al*., 2002; Tiso *et al*., 2002; De Santa Barbara *et al*., 2005; Theodosiou and Tabin, 2005; Davenport *et al*., 2016; Stevens *et al*., 2017) and in hPSC-derived colonic organoids (Munera and Wells, 2017; Múnera *et al*., 2017). In the gastro-duodenal boundary, WNT-mediated crosstalk between the epithelium and mesenchyme are essential for establishing and maintaining a molecular boundary between the gastric and duodenal epithelium (McCracken *et al*., 2017; Kim *et al*., 2005). Evidence from chick studies show that ENCCs regulate the anterior-posterior patterning of stomach (Faure *et al*., 2015), and our data now show that ENCCs express BMP ligands, can posteriorize human gastric epithelium, and that inhibiting BMP signaling prevents the posteriorizing effects of ENCCs.

One striking feature of posteriorized gastric organoids is their ability to form Brunner’s gland-like structures following transplantation and growth *in vivo*. Little is known about the embryonic development of these glands in any species, nor what markers define them. Brunner’s glands normally form in the proximal duodenum close to the pyloric sphincter and lie just below epithelium. We identified a marker profile of human Brunner’s glands; no expression of the gastric markers SOX2, CLD18, MUC5AC, no expression of the duodenal markers CDH17, low expression of CDX2, and positive expression for the gastric mucin MUC5AC and the duodenal marker GLP-1R. Gastric organoids that are mispatterned by ENCCs form a glandular epithelium with a marker profile consistent with Brunner’s glands. Human PSC-derived Brunner’s gland organoids are a new model system to study development of this glandular system and identify the role of ENCCs in patterning the gastro-duodenal region.

Our findings highlight that the only context in which we see formation of normal gastric tissue is when foregut spheroids are combined with a carefully controlled amount of SM and ENCCs. Adding SM alone results in organoids with poorly organized smooth muscle and a simple epithelium and adding ENCCs along results in mispatterned epithelium and the formation of Brunner’s glands. However, recombining robust populations of SM and ENCCs results in well-organized smooth muscle, an organized neuroglial plexus, and the formation of properly patterned gastric glands with chief and parietal cells. Moreover, the neuroglial plexus forms a functional link with the smooth muscle to regulate rhythmic gastric contractions. We conclude that communication between all three germ layers is essential for proper assembly and morphogenesis of stomach tissue.

A possible mechanism to explain why gastric gland morphogenesis requires both ENS and mesenchyme comes from studies of intestinal and lung development. In the intestine of mice and chicks, mesenchymal clusters (Freddo *et al*., 2016; Walton *et al*., 2012) and smooth muscle (Shyer *et al*., 2013) regulate villus morphogenesis and lung branches (Goodwin et al., 2019). Additional studies in chick showed how BMP signaling from both the epithelium and neural cells regulates the radial position of developing smooth muscle layers (Huycke et al., 2019). It follows then that innervation of smooth muscle layers in the human gastric antrum may promote development of organized smooth muscle and glandular morphogenesis. Our data suggest that ENCCs require a robust population of mesenchyme to promote gastric gland morphogenesis. However, addition of ENCCs alone promotes formation of Brunner’s gland-like epithelium in some transplants suggesting that the addition of mesenchyme is important both to maintain gastric identity and/or synergize with the signals coming from the ENCCs. This seems even more plausible when one considers the close proximity and physical connection of enteric nerves with both the smooth muscle and the epithelial cells within stomach glands, both of which are necessary for proper stomach function. In the case of submandibular gland in mouse signals from the ENS maintains the epithelial progenitor pools and supports branching morphogenesis *in vitro* (Knox *et al*., 2010) and *in vivo* (Nedvetsky *et al*., 2014).

Little is known about ENS development and the specifics of neuronal diversity within proximal GI tract, relative to the intestine and colon ENS (Kaelberer *et al*., 2018; Lasrado *et al*., 2017; Rakhilin *et al*., 2016; Bohorquez *et al*., 2015; Walsh and Zemper, 2019; Nagy and Goldstein, 2017; Brookes *et al*., 2013). We have observed differences in ENS development between human gastric and intestinal organoids. For example, in intestinal organoids we did not observe ENCC differentiation into nNOS neurons *in vitro* (Workman *et al*., 2017) whereas these neurons did form in gastric organoids. This suggests that regional differences in ENS development between proximal and distal GI organs could be studied using these human organoid systems. There are many differences in stomach and intestinal development, from orientation and innervation of smooth muscle to glandular morphogenesis and neuronal control of secretion that might be modeled in these systems. Moreover, human organoids are now being used to study human organ physiology. For example, hPSC-derived human intestinal organoids were used as a model of malabsorption in humans and led to the discovery of a new mechanism by which enteroendocrine cells control nutrient absorption (McCauley *et al*., 2020).

In summary, we have generated three germ layer organoids that are morphologically, cellularly, and functionally similar to human stomach tissues. Engineered gastric tissue has glands with surface and pit mucous cells, as wells as chief and parietal cells. We observed oriented layers of smooth muscle that were innervated by functional enteric nerves. We have used this highly manipulable system to begin to define cellular communications that happen during development of the human stomach and expect that it we be equally powerful as a model of gastric diseases. Given that this technology is broadly translatable to other organs, it is possible that engineered tissue might be a source of material for reconstruction of congenital disorders and acute injuries of the upper GI tract.

## LIMITATIONS OF THE STUDY

hPSC-derived gastric organoids can be variable between differentiations. Coordinating the timing of recombination of three simultaneously generated germ layers, as well as the requirement of kidney capsule transplantation into immunocompromised mice necessary to promote three germ layer organoid maturity is technically challenging, lengthy, and can limit protocol scalability.

## ACKNOWLEDGEMENTS

We would like to thank all the members of the Wells, Zorn, and Helmrath laboratories for reagents and feedback. We specifically thank Heather McCauley, Jacob Enriquez, and J. Guillermo Sanchez from the Wells lab for technical assistance with ectopic transplantation surgeries and harvests. We also thank Chris Mayhew and Amy Pitstick from the Pluripotent Stem Cell Facility as well as Matt Kofron and Evan Meyer from the Confocal Imaging Core at Cincinnati Children’s Hospital for constant support and guidance. Finally, we thank Mansa Krishnamurthy, a pediatric fellow in the Wells Lab, for the human antrum and duodenum samples used for Brunner’s Gland characterization. This research was supported by the grants from the NIH, U18 EB021780 (JMW, MAH), U19 AI116491 (JMW), P01 HD093363 (JMW), UG3 DK119982 (JMW), U01 DK103117 (MAH), 1F31DK118823-01 (AKE), NIEHS 5T32-ES007250-29 (DOK), the Shipley Foundation (JMW), and the Allen Foundation (JMW). We also received support from the Digestive Disease Research Center (P30 DK078392).

## AUTHOR CONTRIBUTIONS

AKE and JMW primarily conceived of the experimental design, analyzed the experiments, and co-wrote the manuscript. AKE, DOK, HMB, NS, HMP, LEH, JGS, KK, and MK performed experiments. Specifically, AKE, DOK and HMB advised and performed the organoid recombination experiments. NS, AKE, HMB, JGS performed the ectopic kidney transplantation surgeries and harvests. HMP performed all organ bath experiments. AKE, DOK, and HMB conducted the protein and RNA analysis with LEH greatly serving as technical assistance. LH, JMW, and AMZ conceived of and developed the protocols to direct the differentiation of human PSCs into splanchnic mesenchyme. All authors contributed to the writing and/or editing of the manuscript.

## STAR METHODS

### Animals

All mice used in kidney capsule transplantation experiments were housed in the animal facility at Cincinnati Children’s Hospital Medical Center (CCHMC) in accordance with NIH Guidelines for the Care and Use of Laboratory animals. Animals were maintained on a 12 hour light-dark cycle with access to water and standard chow ad libitum. Healthy male and female immune-deficient NSG (*NOD*.*Cg-Prkdc*^*scid*^*Il2rg*^*tm1Wjl*^*/SzJ*) mice, aged between 8 and 16 weeks old, were used in all experiments. These mice were obtained from the Comprehensive Mouse and Cancer Core Facility. All experiments were performed with the approval of the Institutional Animal Care and Use Committee (IACUC) of CCHMC.

Timed mattings of wildtype mice were used to generate e13.5 embryos for immunohistological analysis. The morning that the vaginal plug was observed was denoted as e0.5.

### Human Biopsy Tissue

The use of human tissues was approved by an Institutional Review Board (IRB) at CCHMC (protocol number 2015-5056). Informed consent for the collection and use of tissues was obtained from all donors, parents, or legal guardians. Full-thickness fundic and antrum stomach tissue samples obtained from bariatric procedures came from the Helmrath Lab at CCHMC under IRB protocol number 2014-0427. Human surgical samples were collected from patients between the ages of 15 and 17, and included both males and females of Caucasian and African American backgrounds. Healthy human full-thickness stomach and duodenal tissue samples were obtained from the CCHMC Pathology Core.

### Human ESC/iPSC lines and maintenance

Human embryonic stem cell (hESC) lines H1 (WA-01) and H9 (WA-09) were purchased from WiCell (NIH approval number NIHhESC-10-0043 and NIHhESC-10-0062). The H1 line is male and the H9 line is female. H9-GAPDH-GFP and H9-GAPDH-mCherry hESCs along with human induced pluripotent stem cell (iPSC) line 77.3-GFP were all generated and obtained from the CCHMC Pluripotent Stem Cell Facility (PSCF) and approved by the institutional review board (IRB) at CCHMC. Human iPSC line WTC11 AAVS1-CAG-GCaMP6f was obtained from Bruce Conklin’s laboratory at UCSF. All hPSC lines were analyzed for pluripotency and the absence of karyotypic abnormalities and mycoplasma contamination by the CCHMC PSCF. Human iPSC line WTC11 was analyzed for karyotype by Cell Line Genetics.

All human hPSCs were maintained in an undifferentiated state as colonies in feeder-free conditions. They were plated on human-ES-cell-qualified Matrigel (BD Biosciences) and maintained at 37°C with 5% CO2 with daily replacement of mTeSR1 media (STEMCELL Technologies). Cells were routinely passaged every 4 days with Dispase (STEMCELL Technologies) after a confluency of about 80-90% was reached.

Differentiations of the following lineages for construction of three-germ layers organoids are not dependent or variable based on starting hPSC line. Please see the Key Resource Table under Experimental Modes: Cell Lines for minimum number of differentiations performed using each cell line.

### Differentiation of hPSCS into splanchnic mesenchyme

Partially confluent hPSCs colonies were dissociated into single cells using Accutase (Thermo Fisher Scientific), resuspended in mTesR1 with thiazovivin (1 μM, Tocris), and passaged 1:20 onto new Geltrex-coated 24-well plates (Sigma Aldrich). The directed differentiation of hPSCs into lateral plate mesoderm has been previously described (Han *et al*., 2020; Loh *et al*., 2016). Briefly, hPSCs were exposed to Activin A (30 ng/ml, Cell Guidance Systems), BMP4 (40 ng/ml, R&D Systems), CHIR99021 (CHIR, 6 μM, ReproCell), FGF2 (20 ng/ml, ThermoFisher Scientific), and PIK90 (100 nM, EMD Millipore) for 24 hours. A basal media composed of Advanced DMEM/F12 (ThermoFisher Scientific) supplemented with B27 supplement (1X, ThermoFisher Scientific), N2 supplement (1X, ThermoFisher Scientific), HEPES (13 mM, ThermoFisher Scientific), L-Glutamine (2 mM ThermoFisher Scientific), and penicillin-streptomycin (1X, ThermoFisher Scientifics) was used for this and all subsequent differentiation steps. Cells were then exposed to A8301 (1 μM, Tocris), BMP4 (30 ng/ml), and C59 (1 μM, Cellagen Technology) for 24 hours. For splanchnic mesoderm generation, cells were cultured in A8301 (1 μM), BMP4 (30 ng/ml), C59 (1 μM), FGF2 (20 ng/ml), and RA (2 μM, Sigma-Aldrich) from Day 2 to Day 4. To further direct regional splanchnic mesoderm, RA (2 μM), PMA (2 μM, Tocris) was used for 2 days, and then RA (2 μM), PMA (2 μM), NOG (100 ng/ml, R&D Systems) was used at the last 1 day to promote esophageal/gastric mesenchyme fate. Medium was changed every day throughout protocol. Confluent cells were resuspended using an Accutase treatment (2–3 min) and immediately combined with hAGOs, hFGOs, and hEOs (see below for recombination procedure).

### Differentiation of hPSCS into ENCCs

The generation of hPSC-derived ENCCs has been previously published (Bajpai *et al*., 2010; Workman *et al*., 2017). Briefly for ENCC generation, confluent hPSCs were treated with collagenase IV (500 U/ml, Gibco) in mTeSR1 for 60-90 mins to detach colonies. Cells were diluted and washed with DMEM/F-12 (Gibco) and then gently triturated and resuspended in neural induction media, 1:1 ratio of DMEM/F12-GlutaMAX (Gibco) and Neurobasal Medium (Gibco) with B27 supplement (0.5x, Gibco), N2 supplement (0.5x, Gibco), pen-strep (1x, Gibco), insulin (5 µg/mL, Sigma-Aldrich), FGF2 (20 ng/mL, R&D Systems), and EGF (20 ng/mL, R&D Systems), on non-TC-treated petri dishes (6cm, Fisherbrand). Neural induction media was changed daily and all-trans RA (2 μM) was added on days 4 and 5 for posteriorization. Day 6 free-floating neurospheres were plated on human fibronectin (HFN, 3 μg/cm^2^, Corning) and fed neural induction media without RA for 4 days. Migrated cells were collected using a 90 sec Accutase treatment and passaged onto HFN. Passaged cells were allowed to grow to confluency for an additional 4 days and fed neural induction media without RA every day. Confluent cells were then collected using a 2-3 min Accutase treatment and immediately combined with hAGOs, hFGOs, and hEOs (see below for recombination procedure).

### Differentiation of hPSCS into hAGOs, hFGOs, and hEOs

We utilized slightly modified previously published protocols to generate hAGOs, hFGOs and hEOs (McCracken et al. 2014, 2017, Trisno et al., 2018). For hAGO and hFGO generation, confluent hPSC cultures were treated with Accutase to resuspend as single cells in mTeSR1 with ROCK inhibitor Y-27632 (10 μM; Tocris) and plated onto a Matrigel-coated 24-well dish (Sigma Aldrich). To direct the differentiation into definitive endoderm (DE), the hPSCs were exposed to Activin A (100 ng/ml) and BMP4 (50 ng/ml) in RPMI 1640 media (Life Technologies). For the following two days, cells were exposed to only Activin A (100 ng/ml) in RPMI 1640 media containing increasing concentrations (0.2% and 2.0%, respectfully) of defined fetal bovine serum (dFBS; HyClone). To then pattern DE into posterior foregut endoderm spheroids, cells were treated with <u>FGF4</u> (500 ng/ml, R&D systems), NOG (200 ng/ml), and CHIR (2 μM) for 3 days, with media changed daily, in RPMI 1640 with 2% dFBS. RA (2 μM) was added on the third day of FGF4/NOG/CHIR treatment.

### Recombination and additional spheroid patterning

Single cell suspensions of mesenchymal cells and ENCCs were counted and added to foregut spheroids at an approximate ratio of 1,000 ENCCs and 2,500 mesenchyme cells per spheroid. Cell mixtures were mixed via gentle pipetting, centrifuged at 300g for 3-5 minutes, and embedded into 50 μL of basement membrane Matrigel to allow three-dimensional *in vitro* culture. Organoids were fed with a base media of Advanced DMEM/F12 supplemented with B27 supplement (1X), N2 supplement (1X), HEPES (13 mM), L-Glutamine (2 mM), penicillin-streptomycin (1X), and EGF (100 ng/mL). In addition to this base media, the first three days were supplemented with NOG (200 ng/mL) and RA (2 μM). In addition to EGF, hFGOs were supplemented with CHIR (2 μM) throughout the organoid outgrowth and also received a 48 hr. pulse of BMP4 (50 ng/mL) and PD0325901 (2 μM, Stem Cell Technologies) 96 hours prior to collection for parietal cell differentiation *in vitro*. Media was replaced every 3-4 days. Two weeks following spheroid embedding in Matrigel, the organoids were collected and re-plated in fresh Matrigel at a dilution of ∼1:12.

### *In vivo* transplantation of hAGOs and hFGOs

hAGO, hFGO, hAGO +ENCC, hAGO +SM, hAGO +SM +ENCC, and hFGO +SM +ENS were all ectopically transplanted into the kidney capsule of NSG mice as previously described (Watson *et al*., 2014). Briefly, four week old hAGOs or hFGOs were removed from Matrigel and transplanted into the kidney subcapsular space. Engrafted organoids were harvested 6–15 weeks after transplantation and analyzed for neuroglial, epithelial, and mesenchymal maturation.

### *Ex vivo* muscle contraction and ENS function

Muscle contraction was assayed as previously described (Poling *et al*., 2018) and ENS function and motility were assayed as previously described with slight modifications (Workman *et al*., 2017). Briefly, strips of tissue approximately 2 x 6 mm in size were dissected and the epithelium mechanically removed in a method similar to seromuscular stripping as previously described (Workman *et al*., 2017). No chelation buffer was used. Resulting strips of muscle from hAGO +SM +ENCC were mounted within an organ bath chamber system (Radnoti) to isometric force transducers (ADInstruments) and contractile activity continuously recorded using LabChart software (ADInstruments). After an equilibrium period, a logarithmic dose response to Carbamyl-β-methylcholine chloride (Bethanechol; Sigma-Aldrich) was obtained through the administration of exponential doses with concentrations of 1 nM to 10 mM at 2 min intervals before the administration of 10 µM scopolamine (Tocris Bioscience). Data are normalized to muscle strip mass. After another equilibrium period, muscle preparations were then stimulated with a control EFS pulse. NG-nitro-L-arginine methyl ester (L-NAME; 50 μM; Sigma) was applied 10 min before EFS stimulation to observe the effects of NOS inhibition. Without washing, Atropine (atropine sulfate salt monohydrate; 1 μM; Sigma) was the applied 10 min prior to a final EFS stimulation to observe the cumulative effect of NOS and Ach receptor inhibition. After several washes and an additional equilibrium period, another control EFS pulse was administered. Neurotoxin tetrodotoxin (TTX; 4 μM; Tocris) was administered 5 min before a final EFS stimulation. Analysis was performed by calculating the integral (expressed as area under the curve, AUC) immediately before and after stimulation for 60s. Data are normalized to muscle strip mass.

### *Ex vivo* GCamP6f calcium imaging

Detection of calcium transients was performed using the above-mentioned human iPSC line WTC11 AAVS1-CAG-GCaMP6f. Transplanted hAGOs +ENCC were harvested and then cultured on 8-well micro-slide (Ibidi) for 24 hours prior to imaging. They were then imaged every 4-15 sec for 3-10 min using either a 10x or 20x objective on a Nikon Ti-E inverted A1 confocal microscope with NIS elements software to obtain background fluorescence level. Transplanted hAGOs +ENCC were then treated with 30 mM KCl. Experiments were carried out at RT.

### Tissue Processing, Immunohistochemistry, and Microscopy

Cell monolayers, ENCCs, and day 0 spheroids were washed with 1x phosphate-buffered saline (PBS), fixed with 4% paraformaldehyde (PFA) at room temperature (RT) for 15 min, washed, and stored in PBS at 4°C. Four week old *in vitro* organoids and *in vivo* transplants were washed with PBS, fixed in 4% PFA at 4°C overnight, washed, and then placed in either PBS, 30% sucrose in PBS, or 70% ethanol at 4°C overnight for downstream whole mount, cryogenic, or paraffin processing, respectively. Prior to fixation, whole mount tissues were extracted from Matrigel using manual pipetting in cold PBS and Cell Recovery Solution (Corning). Tissues were then embedded in either O.C.T. Compound (Tissue-Tek) or paraffin and were serially sectioned at a thickness of 7-8 μm onto Superfrost Plus glass slides (Fisherbrand). Cryosection slides and paraffin slides were stored at -80°C and RT, respectively. Routine Hematoxylin & Eosin (H&E) staining was performed by the Research Pathology Core at CCHMC.

Frozen slides were thawed to room temperature (RT) and rehydrated in PBS, while paraffin slides were deparaffinized, rehydrated, and subjected to heat-and pressure-induced antigen retrieval in citrate buffer (0.192% citric acid and 0.0005% Tween 20 in dH_2_0 of pH 6.0 with NaOH) for 30 minutes and brought to RT on ice. All slides and cells were washed with PBS, permeabilized with 0.5% Triton X-100 in PBS (PBST) for 15 min at RT and then blocked with 5% normal donkey serum (NDS, Jackson ImmunoResearch) in PBS for one hour at RT. Tissue was incubated at 4°C overnight in primary antibodies diluted in 5% NDS in PBS. Specific antibody details are listed in the Key Resource Table. The following day, tissue was washed and incubated with secondary antibodies at RT for one hour, thoroughly washed, and cover slipped with Fluoromount-G (Southern Biotech).

For wholemount staining, organoids were washed at RT and then permeabilized with PBST at 4°C overnight. The next day, organoids were blocked in 5% NDS in PBST for 6-8 hours at RT and then incubated in primary antibodies at 4°C overnight on a rocking platform. Organoids were extensively washed in PBST and then incubated in secondary antibodies at 4°C overnight. Finally, organoids were washed with PBST, PBS and then serially dehydrated to 100% methanol. Organoids were then optically cleared with Murray’s Clear (2:1 benzyl benzoate: benzyl alcohol, Sigma) for at least 15 minutes prior to imaging.

Brightfield and GFP fluorescence images of live tissue samples were captured using either a Leica DMC5400 or DFC310 FX camera attached to a stereomicroscope. Whole mount and all immunofluorescent images were captured using a Nikon Ti-E inverted A1 confocal microscope. Images were processed and quantified using Nikon NIS Elements, Bitplane Imaris, Adobe Illustrator, and Microsoft PowerPoint software.

### RNA isolation and quantitative real-time PCR (qRT-PCR)

Spheroids and organoids were harvested in RA1 Lysis Buffer and β-mercapethanol and stored at -80°C until total RNA was isolated using NucleoSpin RNA Isolation Kit (Macherey-Nagel) according to manufacturers’ instructions. Complementary DNA (cDNA) was reverse transcribed from 116 ng of RNA using a SuperScript VILO cDNA Synthesis Kit (Invitrogen). qRT-PCR was performed using a QuantiTect SYBR Green PCR Kit (Qiagen) in MicroAmp EnduraPlate Optical 96-Well Fast Reaction Plates (Applied Biosystems) and run on a QuantStudio 6 Real-Time PCR Detection System (Applied Biosystems). Primer sequences are listed in the Key Resource Table. Analysis was performed using the ΔΔCt method by first normalizing all cycle threshold (Ct) values to a base housekeeping gene (*GAPDH, PPIA, or FOXF1*) and then to the control hAGO samples. Statistical analysis was performed using Student’s *t*-test.

### Statistical analyses

For analysis of organoid patterning, “n” represents the number of replicates performed in each experiment and each replicate is defined as 1 well of approx. 3-5 organoids in Matrigel culture. Any analysis presented from one individual experiment is representative of trends seen across at least two individually seeded experiments. All data are represented as mean ± s.d. Student’s *t*-tests with 2-tailed distribution and un-equal variance was completed using Microsoft Excel, where p ≤ 0.05 is symbolized by *, p ≤ 0.01 is symbolized by **, and p ≤ 0.001 is symbolized by ***. The determined significance cutoff was p ≤ 0.05. No statistical method was used to predetermine sample size. The investigators were not blinded to allocation during experiments and outcome assessment. No randomization was made.

## SUPPLEMENTAL INFORMATION

**Figure S1**. Splanchnic mesenchymal recombination yielded the most added exogenous mesenchyme while still retaining endogenous mesenchyme, relating to Figure 1. (A) Brightfield images of 4 week *in vitro* hAGOs recombined with varying concentrations of splanchnic and septum transversum (STM) mesenchyme on day 6 of hAGO protocol. Visual qualitative assessment of 4 week *in vitro* hAGOs lead to utilizing splanchnic mesenchyme at a ratio of 50,000 cells/well of approx. 20-30 hAGO spheroids. This equates to an approx. 2:1 ratio of splanchnic mesenchymal cells to hAGO epithelial cells. (B) Brightfield images of hAGOs grown for four weeks *in vitro* with and without recombination with exogenous GFP-labeled gastric-esophageal mesenchyme (GEM) (green) costained with mesenchymal marker FOXF1 (red). Higher magnification images are shown to the right. This relates to Fig. 1D. (C) Quantification of various mesenchymal recombination techniques, including day 6 mesenchymal recombination (left) of either GFP+ splanchnic (SM) or cardiac (CM) mesenchyme and day 9 mesenchymal recombination (right) of either GFP+ gastric/esophageal (GEM) or septum transversum (STM) mesenchyme (n=at least 6 fields from at least 3 organoids per condition from one differentiation, same trend seen across at least two individually seeded differentiations, **p<0.01, Student’s t-test).

**Figure S2**. Three germ layer *in vitro* and *in vivo* hAGOs and hFGOs contain GFP+ splanchnic mesenchyme and RFP+ ENS, relating to Figure 3. (**A**) Brightfield and fluorescent images of four week *in vitro* hAGO +GFP SM (green +RFP ENS (red) and epithelial ECAD (white). Higher magnification images are show on the bottom row. (**B**) Quantification of GFP+ mesenchyme, RFP+ neural, and ECAD+ epithelial populations within four week *in vitro* hAGOs (n=8 fields from at least 3 organoids from one differentiation, same trend seen across at least two individually seeded differentiations). (**C**) Representative images of gross *in vitro* and post transplantation hAGOs with and without incorporation of SM and GFP-labeled ENS. GFP neurons formed networks around grafts post transplantations. (**D**) Quantification of mesenchymal populations within four week *in vitro* hAGOs (n=11-18 fields from at least 3 organoids from one differentiation, same trend seen across at least two individually seeded differentiations, ***p<0.001, Student’s t-test). (**E**) Representative images and (**F**) quantification of the epithelial (1), proximal muscularis mucosa (2), submucosa (3), and distal muscularis externa (4) layer thickness from hAGOs 12 weeks post transplantation, 38 week old human fetal stomach, and adult stomach (n=3-9 fields from 3 hAGOs, 1 38wk fetal stomach, and 1 adult stomach). (**G**) Representative images of gross *in vitro* and post transplantation hFGOs with and without incorporation of SM and GFP-labeled ENS. GFP neurons formed networks around grafts post transplantations (**H**) Representative histological (left) and immunofluorescent (middle and right) comparison of *in vitro* hFGOs with and without added SM and GFP ENS as well as with and without added BMP4 and MEK pathway inhibitor PD03 to stimulate parietal cell differentiation. Neurons are labeled with TUJ1 (green, middle), smooth muscle with αSMA (red, middle), and parietal cells with APT4A (green, right) and ATP4B (red, right). Epithelium is labeled with ECAD (white). Inset (right) highlighting ATP4B+ parietal cell differentiation in hFGOs with added SM and ENS.

**Figure S3**. Constructing three germ layer organoids *in vitro* is applicable to human esophageal organoids, relating to Figure 3. (**A**) Brightfield and GFP-fluorescent images of 1 mo. *in vitro* HEOs. GFP cells label exogenous hPSC-derived SM. Immunofluorescent images of representative HEOs depicting GFP+ (green), FOXF1+ mesenchymal (red), and Vimentin+ (VIM, red) mesenchymal cells. (**B**) Quantification of different mesenchymal populations within 1 mo. *in vitro* HEOs. FOXF1+ expressing cells without GFP mark endogenous mesenchyme, while both GFP+ groups represent exogenous SM (n=16-18 fields from at least 3 organoids per condition from one differentiation, same trend seen across at least two individually seeded differentiations, **p<0.01, ***p<0.001, Student’s t-test). (**C**) Brightfield images of 2 mo. *in vitro* HEOs +/-ENCC showing a visible expansion of additional cells within HEOs incorporated with ENS. (**D**) Immunofluorescent images of 1 mo. *in vitro* HEOs depicting TUJ1+ (green) enteric neurons surrounding the KRT5+ (red) and ECAD+ (white) epithelium of HEOs +ENCC. Higher magnification images are shown to the right. (**E**) Relative expression of neuronal-specific genes including tubulin genes, TUJ1 and MAP2, and filament genes, Nestin within 1 mo. HEOs +ENCC (n=3, representative of 3 individual experiments, **p<0.01, ***p<0.001, Student’s t-test). (**F**) Brightfield and fluorescent images of 1 mo. *in vitro* HEOs +GFP SM +RFP ENCC. Higher magnification images are show on the bottom row. ECAD marks the epithelium in white. (**G**) Quantification of GFP mesenchyme, RFP neural, and ECAD+ epithelial populations within 1 mo. *in vitro* HEOs (n=12 fields from at least 3 organoids from one differentiation, same trend seen across at least two individually seeded differentiations). (**F**) Human tissue sample of 38 week esophagus (H&E, top; TUJ1, green, bottom; αSMA, red, bottom; ECAD, white; bottom; DAPI, blue; bottom). (I Brightfield images of 1 mo. *in vitro* HEOs +/-SM +/-ENCC showing a visible expansion of additional cells within HEOs incorporated with ENS. Immunofluorescent images of 1 mo. *in vitro* HEOs depicting TUJ1+ (green) enteric neurons and FOXF1+ (red) mesenchyme surrounding the KRT8+ (white) epithelium of HEOs +SM +ENS. Higher magnification images are shown to the right.

**Figure S4**. hPSC-derived ENCCs differentiated into neuroglial subtypes when engineered into hAGOs without exogenous mesenchyme, relating to Figure 5. (**A**) Schematic depicting detailed method of deriving and innervating hAGOs. (**B**) Representative brightfield (left) and GFP fluorescent (right) images of four week *in vitro* hAGOs with and without GFP+ ENS. (**C**) Representative images of end time point, day 14, monolayer ENCCs stained for key ENCC markers SOX10 (green, left), AP2A (red, middle), and p75 (red, right). (**D**) Relative expression of neural crest specifier genes (SOX9, SOX10, and SNAIL2), and (**E**) regional hox patterning genes (HOXB3, HOXB5, HOXB7) (n=3 wells from one differentiation, same trend seen across at least four individually seeded differentiations, **p<0.01, Student’s t-test). (**F**) Wholemount immunofluorescence of four week *in vitro* hAGO +ENCC labeled with TUJ1+ neurons. (**G**) Immunofluorescent images of TUJ1+ neurons (top) and S100b+ glial cells (bottom) co-expressed with GFP labeled ENCCs. (**H**) Quantification of the neuroglial composition co-expressing GFP (n=6 fields from one differentiation, same trend seen across at least two individually seeded differentiations, ***p<0.001, Student’s t-test). (**I**) Immunofluorescent images of specific neuronal subtypes, including inhibitory neurons (nNOS) and synaptophysin (SYNAP), dopaminergic neurons (TH), sensory neurons (CALB1), and glial fibrillary acidic protein (GFAP) in hAGOs +ENCC. (**J**) Quantification of GFAP+ cells in four week *in vitro* hAGO +ENCC (n=11-16 fields from at least 3 organoids from one differentiation, p-value determined using Student’s t-test). Representative images (**K**,**M**) and quantification (**L**,**N**) of (**K**,**L**) TUJ1+ neurons (red) and (**M**,**N**) nNOS+ inhibitory neurons (green) within four week *in vitro* hAGOs +ENCC (top) and e13.5 WT murine stomach (bottom) (n>2 fields from one differentiation and one mouse; there is no significant difference). Epithelium is labelled with ECAD (white). Right panels are higher magnification insets of left panels.

**Figure S5**. ENS cells support *in vivo* growth and survival of hAGOs, relating to Figure 6. (**A**) Schematic illustration the method of transplanting hAGOs +ENCC. (**B**) Quantification of epithelial growth from transplanted hAGOs with and without ENS (n=46-48 transplants per condition from 6 individual differentiations). (**C**) Representative brightfield (left) and GFP fluorescent (right) images of transplanted hAGOs with and without GFP+ ENS following *in vivo* transplantation (n=29). (**D**) Brightfield (left) and immunofluorescent (right) images of ECAD+ epithelium (white) from *in vivo* hAGOs with or without ENS cystic grafts. Representative images of (**E**) differentiated antral epithelial and (**F**) mesenchymal and neuronal cell types in hAGOs +ENCC following *in vivo* growth. (**E**) Endocrine cells (yellow arrow) are marked with gastrin, ghrelin, somatostatin, and serotonin, as well as surface mucous cells marked by MUC5AC. (**F**) Mesenchymal cells are marked with FOXF1+ with smooth muscle marked with αSMA. Lineage-traced hPSC-derived ENCCs are marked by GFP and differentiated inhibitory neurons are marked with nNOS. Sections were counterstained with epithelial marker ECAD (white) and nuclear DAPI (blue).

**Figure S6**. Transplanted hAGO grafts +ENS contain appropriate neuroglial cell types that are able to efflux calcium, relating to Figure 6. (**A**) Immunofluorescent images of *in vivo* hAGOs +ENCC show presence of TUJ1+ neural and S100b+ glial cells as well as differentiated neuronal subtypes marked by peripherin and nNOS. ECAD (white) marks the epithelium. (**B**) Immunofluorescent images of *in vivo* hAGOs +ENCC show presence of GFAP+ glial cells (green). (**C**) Quantification of GFAP+ cells in 14 week *in vivo* hAGO +ENCC (n=10 fields from at least 3 organoids from one differentiation). (**D**) Wholemount immunohistochemistry of *in vivo* hAGO +ENCC show a 3D network formation of TUJ1+ neurons within αSMA+ smooth muscle layers. (**E**) Brightfield images of the *in vivo* hAGO +ENCC grafts used to obtain live images of GCaMP neuronal firing. (**F**) GFP fluorescent static images taken from live-imaged movie depicting firing of two individual neurons, indicated by a yellow arrow and orange arrowhead.

**Figure S7**. Defining Brunner’s gland epithelium using combinatorial marker expression analysis of Human Brunner’s Glands, relating to Figure 7. (**A**) H&E (left) and immunofluorescent (right) images of adult human Brunner’s glands labeled with intestinal epithelial marker CDH17 (white). (**B**) Immunofluorescent comparison of adult human antrum (top) and duodenum and Brunner’s glands (bottom). The gastric epithelial cell types are labeled with CLDN18, SOX2, MUC5AC, PGA3 (red), and PGC (green, right). Intestinal cell types are labeled with markers CDX2, and MUC2 (green). Endocrine hormone GAST (green, middle left) was observed in all regions. Only PGC and GAST (green) were consistently observed in Brunner’s Glands. Epithelium was labeled with ECAD or β-catenin (white).

**Figure S8**. Brunner’s Gland-like epithelium only developments from hAGOs innervated by ENCCs untreated with Noggin and Retinoic Acid, relating to Figure 7. (**A**) Relative expression of BMP ligands (*BMP4* and *BMP7*) at different points of ENCC differentiation. hPSCs and day 6 neurospheres (NSs) were used to compare to ENCCs. (**B**) Relative expression of BMP ligands with and without NOG and RA treatment. (**C**) Representative images of organoids with ECAD+ epithelium (white) from transplanted hAGOs +ENCC following recombination at either day 6 or day 9 of hAGO protocol. (D) Representative images of organoids with ECAD+ epithelium (white) and human nuclei expression (green) from transplanted hAGOs +ENCC at day 9 of hAGO protocol; higher magnification is shown to the right.

**Movie S1**. 3-dimensional video image of wholemount immunofluorescence of 10 week *in vivo* three germ layer hAGO +SM +GFP ENS (neurons-green, αSMA-red, ECAD-white, DAPI-blue, 10x magnification). Video corresponds to Figure 2A, bottom right panel.

**Movie S2**. 3-dimensional video image of wholemount immunofluorescence of 4 week *in vitro* hAGO +GFP ENS demonstrates network morphology of TUJ1+ neurons (neurons-green, TUJ1-red, ECAD-white, DAPI-blue, 10x magnification). Video corresponds to Figure S3C, relating to Figure 5.

**Movie S3**. 3-dimensional video image of wholemount immunofluorescence of 8 week *in vivo* hAGO graft +GFP ENS demonstrates network morphology of TUJ1+ neurons (neurons-green, TUJ1-red, ECAD-white, DAPI-blue, 10x magnification). Video corresponds to Figure S5B, relating to Figure 6.

**Movie S4**. Time-lapse video of live-imaged *in vivo* hAGO graft +ENS where ENS was derived from ENCCs containing a GCaMP6f reporter. GFP fluorescent firing demonstrates Ca2+ flux of multiple individual neurons. hAGOs were generated with H1 cells, which do not have a Ca2+ indicator. Images were collected every 4 seconds for 3 minutes using a 20x objective. Video corresponds to Figure S5D, relating to Figure 6.

**Table S1**. Additional brightfield images of *in vivo* hAGO grafts recombined with either mesenchyme, ENCCs, neither, or both, relating to Figure 2.

**Table S2**. List of all neural markers assessed within *in vitro* and *in vivo* organoid cultures, relating to Figure 5.

