## Supplemental Figures for "Engineering functional human gastrointestinal organoid tissues using the three primary germ layers separately derived from pluripotent stem cells"

**Figure S1.** Splanchnic mesenchymal recombination yielded the most added exogenous mesenchyme while still retaining endogenous mesenchyme, relating to Figure 1.

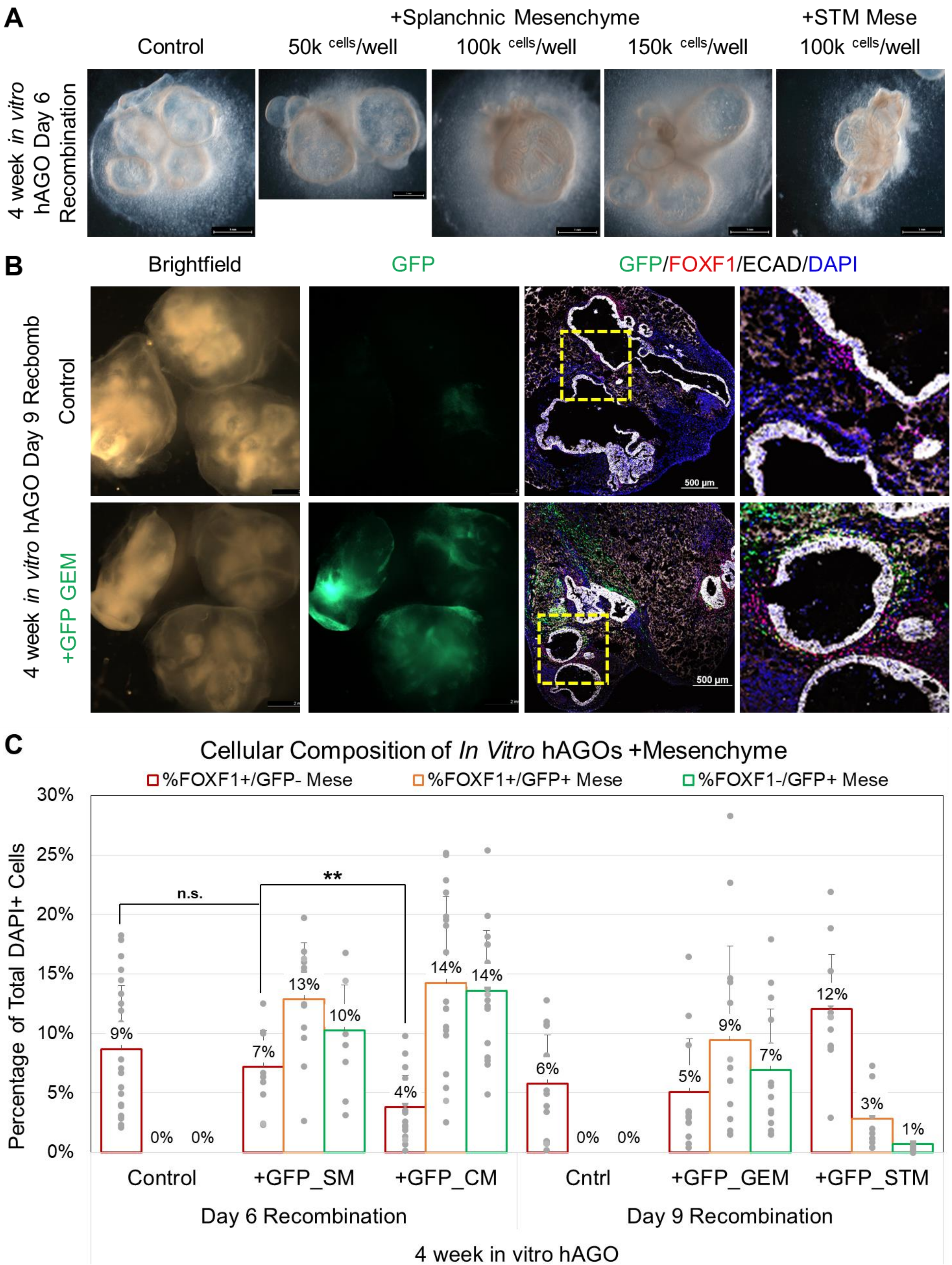

**Figure S2.** Three germ layer *in vitro* and *in vivo* hAGOs and hFGOs contain GFP+ splanchnic mesenchyme and RFP+ ENS, relating to Figure 3.

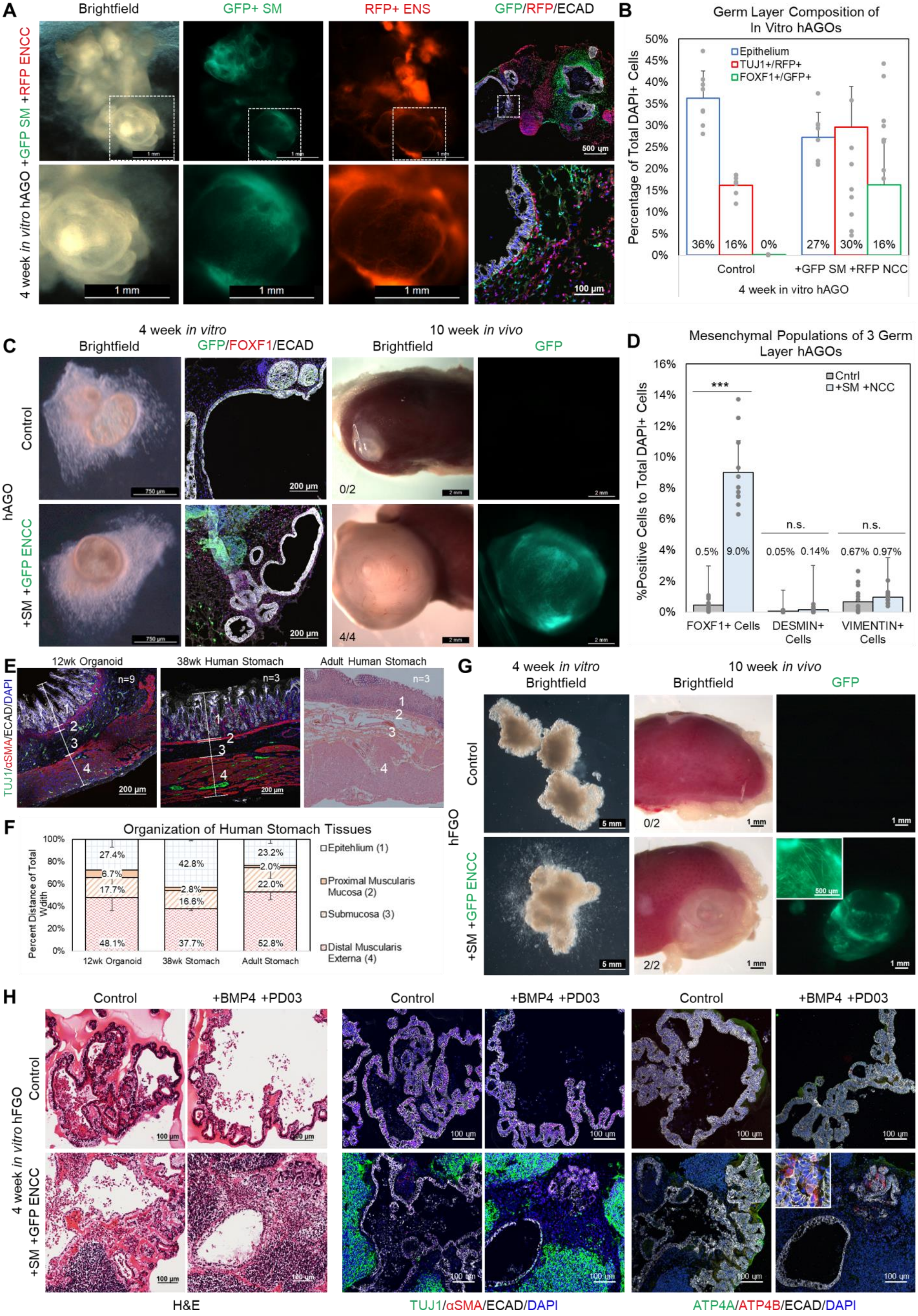

**Figure S3.** Constructing three germ layer organoids *in vitro* is applicable to human esophageal organoids, relating to Figure 3.

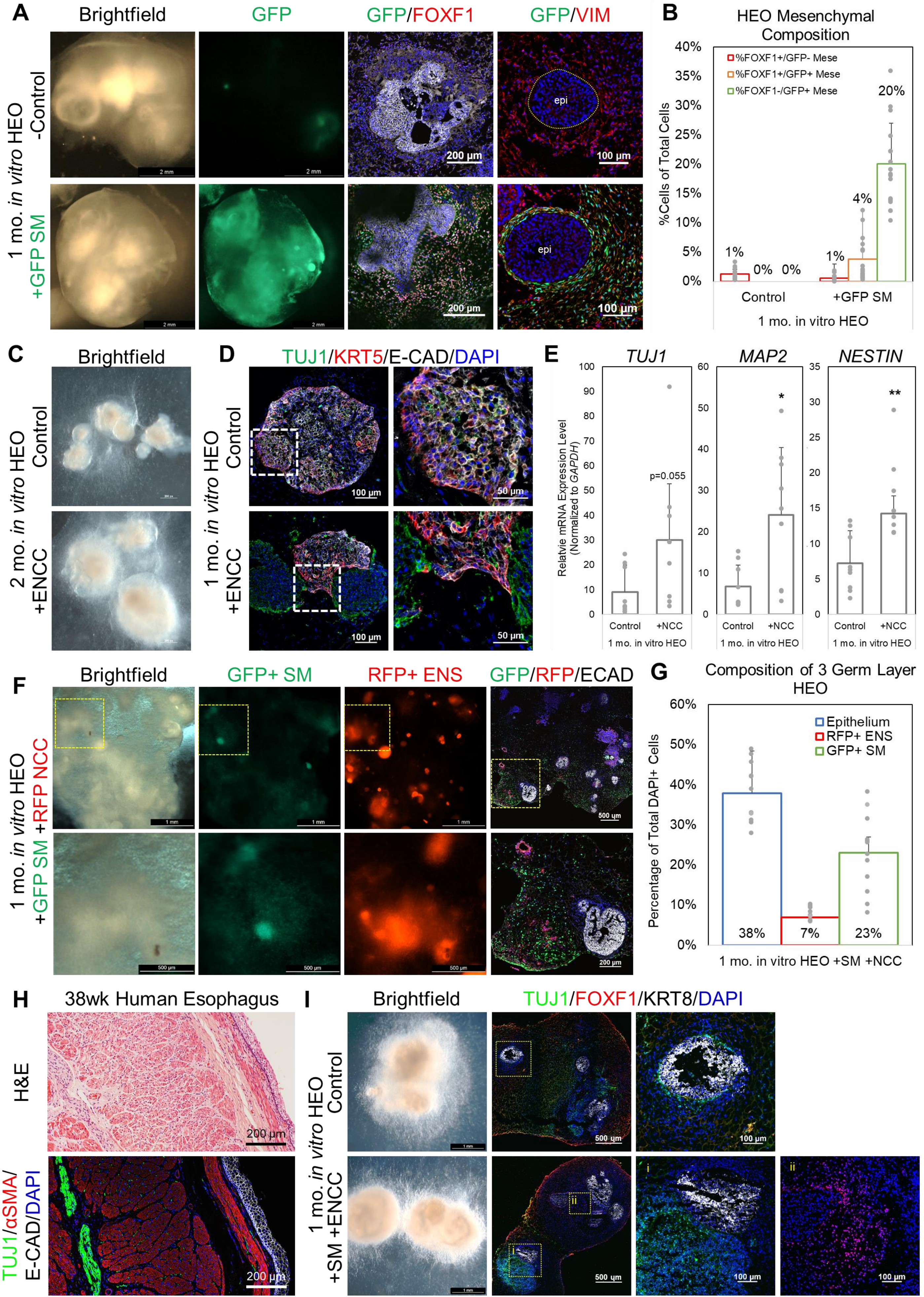

**Figure S4.** hPSC-derived ENCCs differentiated into neuroglial subtypes when engineered into hAGOs without exogenous mesenchyme, relating to Figure 5.

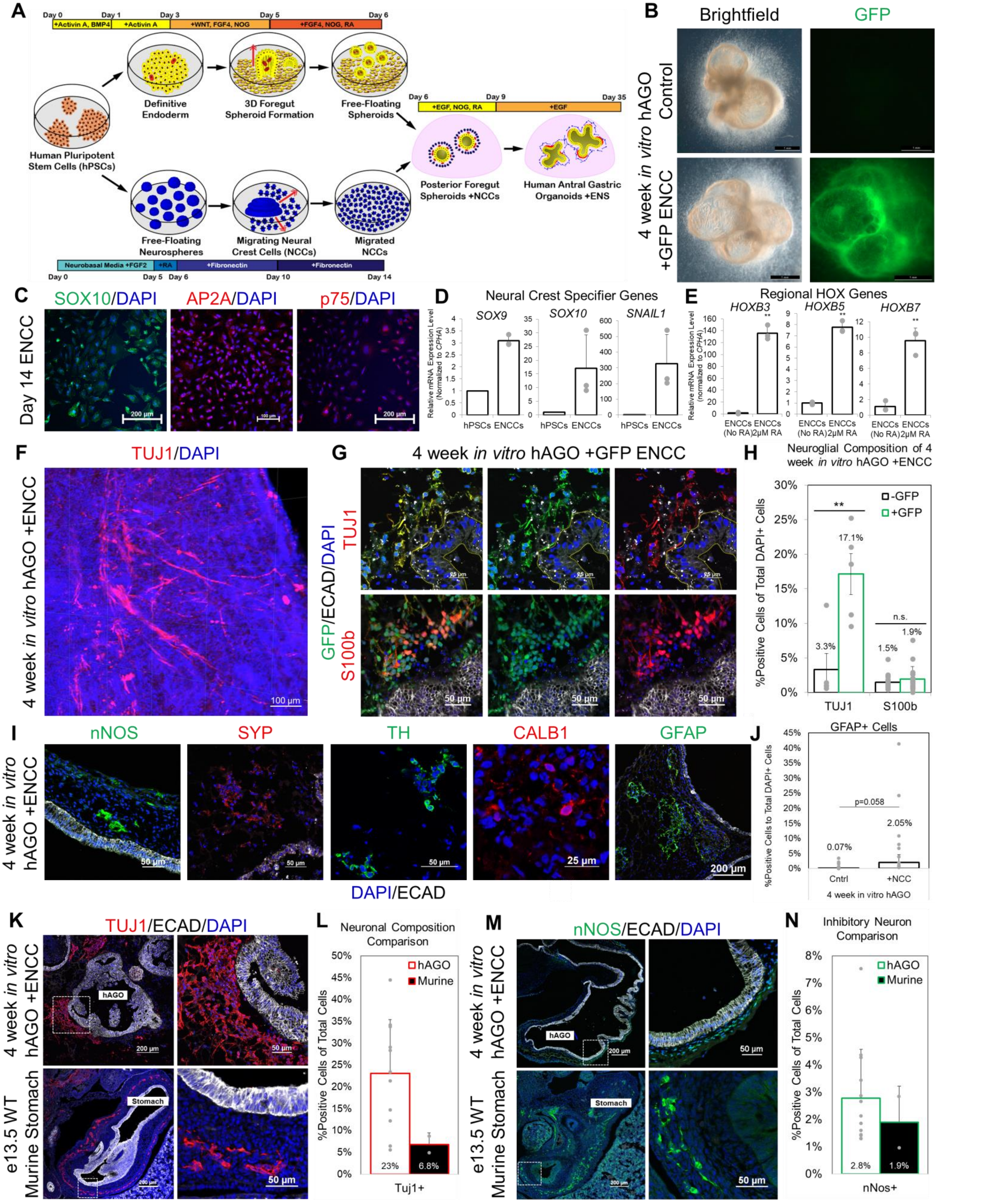

**Figure S5.** ENS cells support *in vivo* growth and survival of hAGOs, relating to Figure 6.

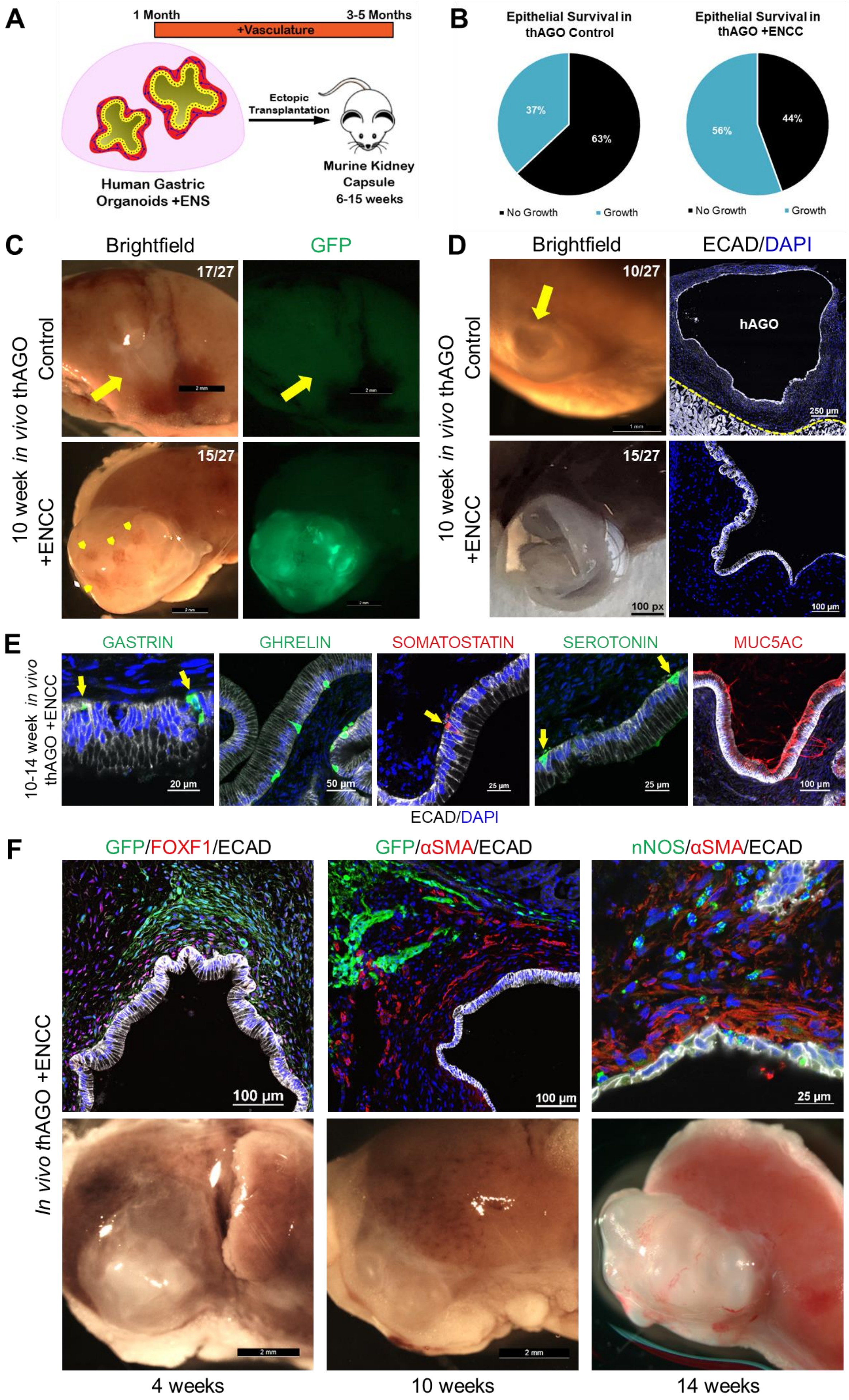

**Figure S6.** Transplanted hAGO grafts +ENS contain appropriate neuroglial cell types that are able to efflux calcium, relating to Figure 6.

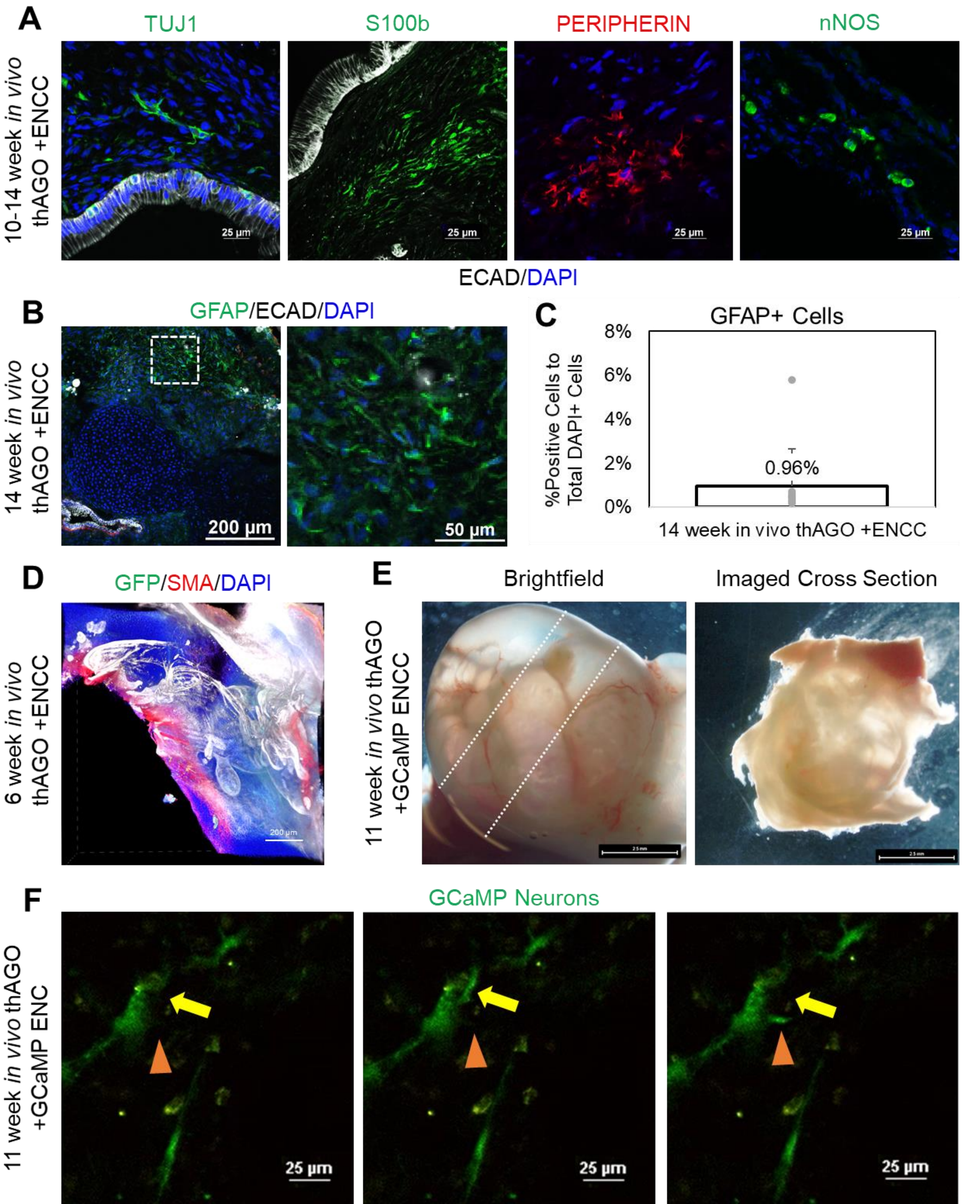

**Figure S7.** Defining Brunner's gland epithelium using combinatorial marker expression analysis of Human Brunner's Glands, relating to Figure 7.

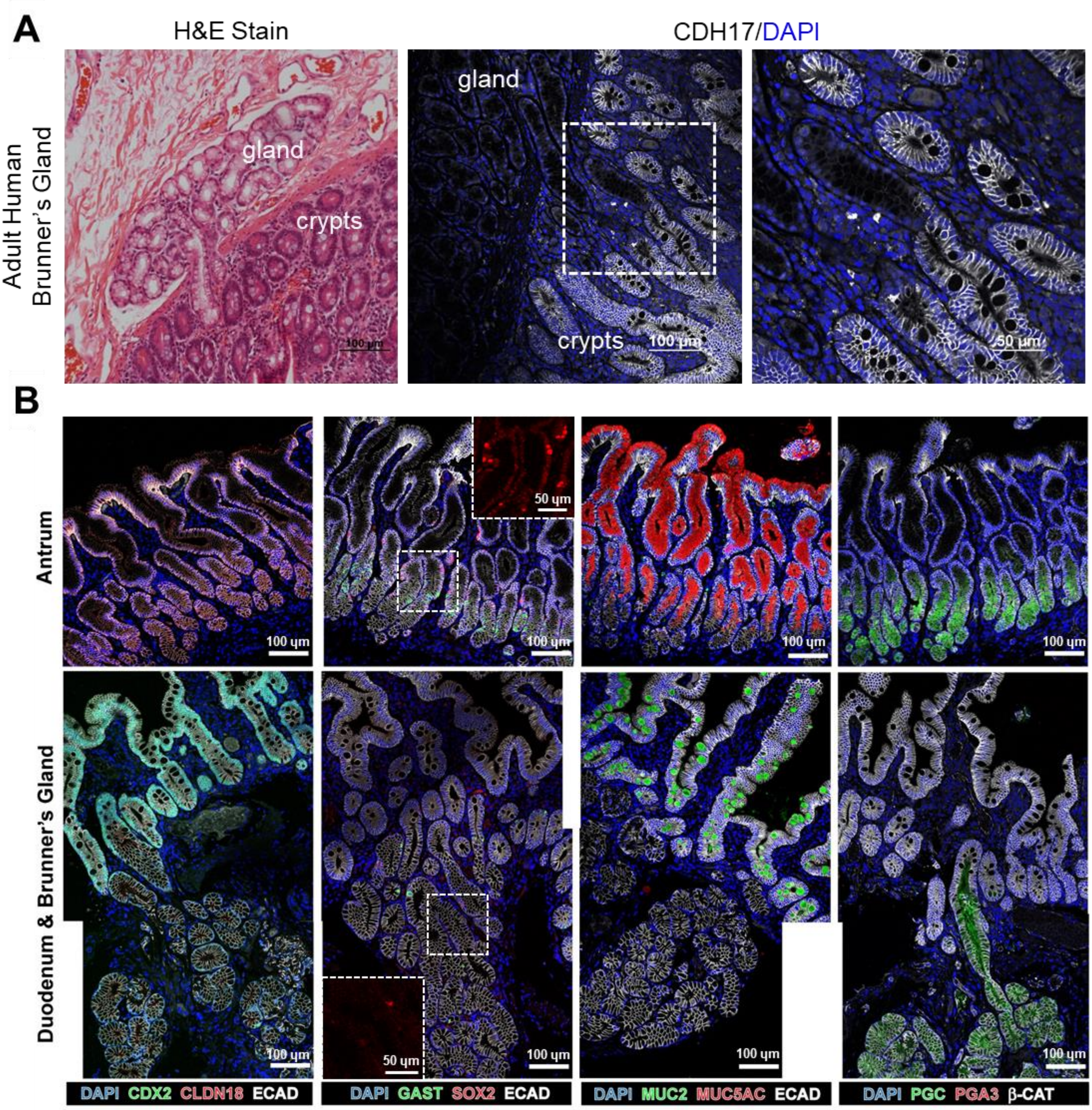

**Figure S8.** Brunner's Gland-like epithelium only developments from hAGOs innervated by ENCCs untreated with Noggin and Retinoic Acid, relating to Figure 7.

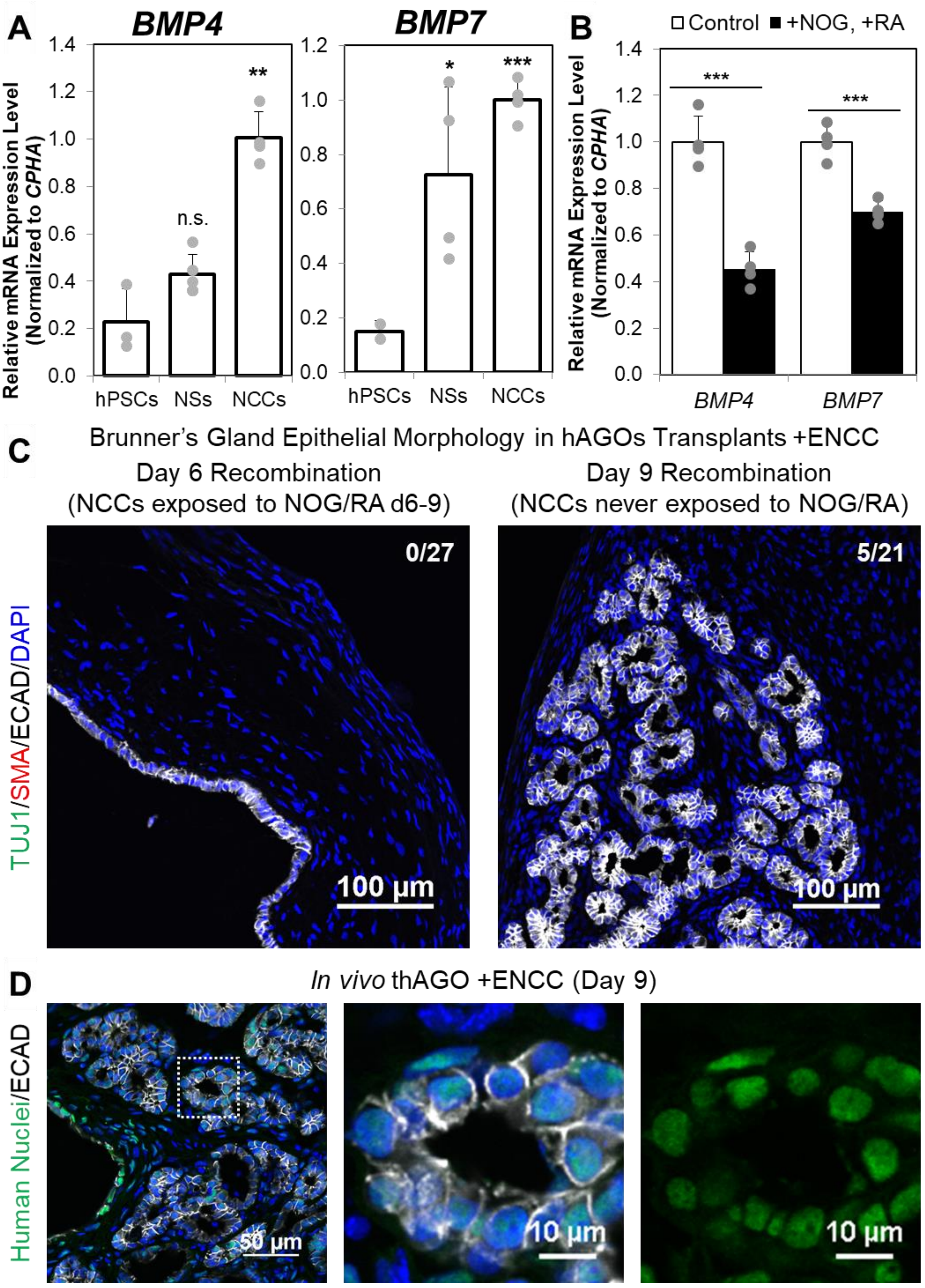
